# mRNA interactions with disordered regions control protein activity

**DOI:** 10.1101/2023.02.18.529068

**Authors:** Yang Luo, Supriya Pratihar, Ellen H. Horste, Sibylle Mitschka, Antonia S. J. S. Mey, Hashim M. Al-Hashimi, Christine Mayr

## Abstract

The cytoplasm is compartmentalized into different translation environments. mRNAs use their 3′UTRs to localize to distinct cytoplasmic compartments, including TIS granules (TGs). Many transcription factors, including MYC, are translated in TGs. It was shown that translation of proteins in TGs enables the formation of protein complexes that cannot be established when these proteins are translated in the cytosol, but the mechanism is poorly understood. Here we show that MYC protein complexes that involve binding to the intrinsically disordered region (IDR) of MYC are only formed when MYC is translated in TGs. TG-dependent protein complexes require TG-enriched mRNAs for assembly. These mRNAs bind to a new and widespread RNA-binding domain in neutral or negatively charged IDRs in several transcription factors, including MYC. RNA-IDR interaction changes the conformational ensemble of the IDR, enabling the formation of MYC protein complexes that act in the nucleus and control functions that cannot be accomplished by cytosolically-translated MYC. We propose that certain mRNAs have IDR chaperone activity as they control IDR conformations. In addition to post-translational modifications, we found a novel mode of protein activity regulation. Since RNA-IDR interactions are prevalent, we suggest that mRNA-dependent control of protein functional states is widespread.

## Introduction

The cytoplasm is compartmentalized by several translation-competent condensates, including TIS granules (TGs)^1-6^. TGs are generated through assembly of the RNA-binding protein TIS11B together with its bound mRNAs^1^. mRNAs whose 3′UTRs are predominantly bound by TIS11B localize to TG^1,6^. Translation in TIS granules allows proteins to form specific protein complexes that cannot be established upon translation outside of TGs^1^ through mechanisms that are not understood.

To comprehensively identify TG-translated proteins, we used fluorescent particle sorting and determined the mRNAs enriched in TGs compared with the cytosol^6^. TG-enriched mRNAs mostly encode low-abundance proteins with a substantial overrepresentation of transcription factors, including MYC^6^. MYC controls different transcriptional programs to regulate a large number of cellular processes, including proliferation, apoptosis, and metabolism^7^. The diverse functions of MYC are mediated by at least 80 different protein interactors^8^. Here, we studied the behavior of MYC to gain insights into the mechanisms controlling TG-dependent protein functional states.

We investigated whether the location of MYC translation within the cytoplasm influences the formation of MYC protein complexes. We observed that MYC complexes established upon MYC translation in TGs involve binding to the MYC IDR whereas those formed in the cytosol involved binding to the folded protein domain. Mechanistically, we found that TG-dependent MYC complexes require TG-enriched mRNAs for their formation. The binding of mRNA to the MYC IDR is not electrostatic but is mediated by a newly discovered RNA-binding domain in IDRs that consists of an α-helix in a serine-rich sequence context and is found in thousands of proteins. Through NMR spectroscopy, we observed that mRNA interaction with the MYC IDR changed the IDR conformational ensemble, resulting in change of chemical shifts. Our results suggest that mRNA-IDR interactions are a widespread mechanism to control protein complex assembly and the activity of proteins with IDRs.

## Results

We showed previously that mRNAs with 3′UTR-bound TIS11B localize to TIS granules, whereas deletion of these 3′UTRs results in cytosolic mRNA localization^1,6^. Using the *MYC* 3′UTR to control *MYC* mRNA localization, we tested whether MYC protein complex assembly is controlled by the location of MYC translation. cDNA expression constructs that contain the MYC coding region together with its 3′UTR (MYC-U) generate *MYC* mRNA transcripts that localize to TGs, whereas omission of the 3′UTR in the constructs (MYC-NU, no UTR) results in cytosolic mRNA localization (Fig. 1a, ED Fig. 1a-d). mRNA localization to TGs or the cytosol is also controlled in a 3′UTR-dependent manner for *SNIP1* which encodes another TG-translated transcription factor (ED Fig. 1d-f).

**Figure 1.**
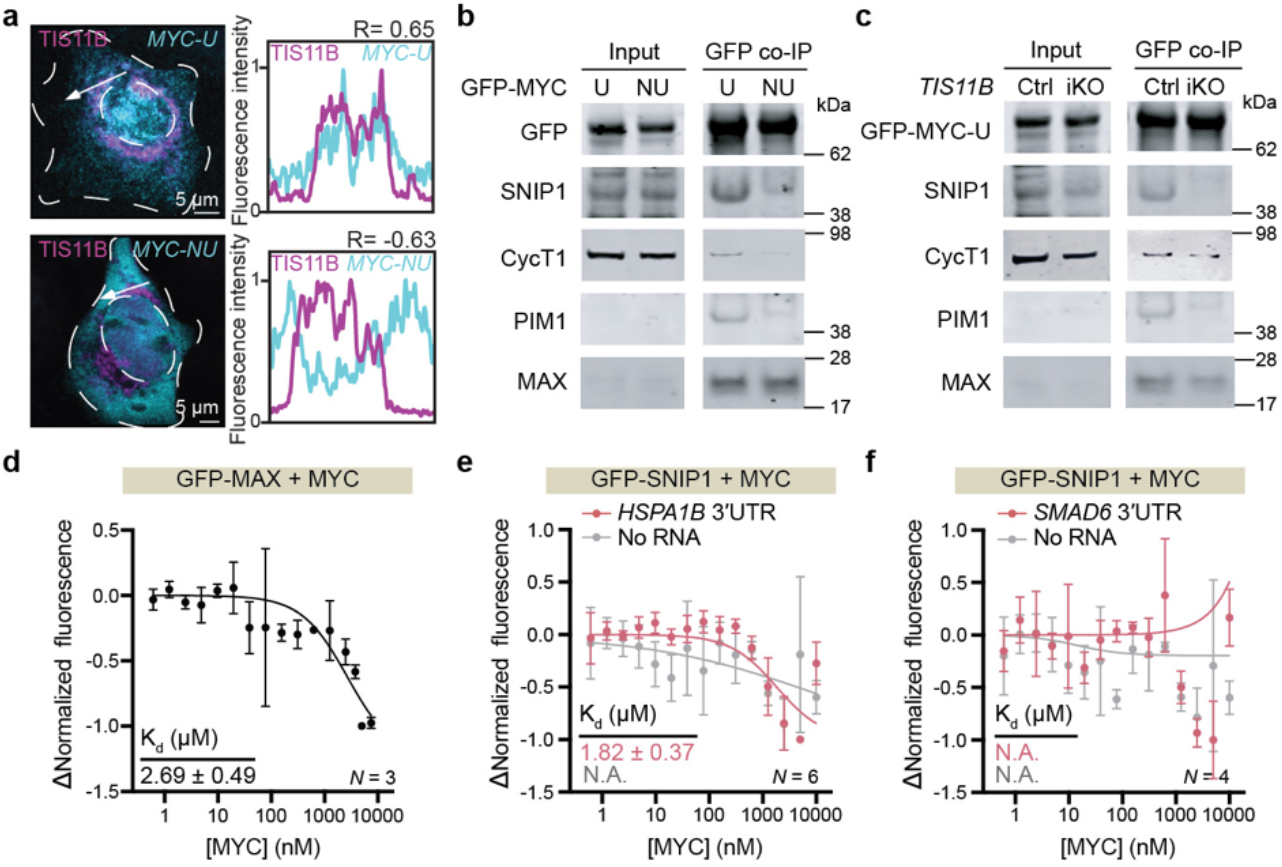
RNA is required for TIS granule-dependent MYC protein complex assembly. **a**, *MYC* mRNA localizes to TGs in a 3′UTR-dependent manner. RNA-FISH (cyan) against GFP after transfection of a cDNA containing GFP-MYC with its 3′UTR (*GFP-MYC-U*, top) or without its 3′UTR (*GFP-MYC-NU*, bottom) in HeLa cells. BFP-TIS11B (magenta) was co-transfected. Right panel, line profiles of fluorescence intensities including Pearson’s correlation coefficients (R). Representative images are shown. Quantification of additional cells is shown in ED Fig. 1d. **b**, Co-IP of endogenous SNIP1, CycT1, PIM1 and MAX using GFP-Trap after transfection of HeLa cells with *GFP-MYC-U* or *GFP-MYC-NU*. 2.5% of input was used. **c**, GFP co-IP of endogenous MYC interactors following transfection of *GFP-MYC-U* into doxycycline-induced *TIS11B* knockout (iKO) versus control (Ctrl) HeLa cells containing non-targeting guide RNAs. **d**, Mean ± std of MST measurement of the MAX-MYC interaction *in vitro*. **e**, Mean ± std of MST measurement of the SNIP1-MYC interaction *in vitro* in the absence or presence of 200 nM *HSPA1B* 3′UTR. N.A., not applicable, as a sigmoidal binding curve cannot be generated. **f**, As in **e**, but 200 nM of *SMAD6* 3′UTR was used.

Previously performed cytoplasmic fractionation revealed that endogenous *MYC* mRNA and several of the known MYC protein interactors have a biased subcytoplasmic mRNA localization pattern and are enriched in TGs compared with the cytosol (ED Fig. 1g, Supplementary Table 1)^6,8^. We hypothesized that interaction partners encoded by TG-biased mRNAs interact with MYC protein in a 3′UTR-dependent manner. Among the known MYC interactors, we set out to test 3′UTR-dependent MYC binding to SNIP1, Cyclin-T1 (CycT1), and PIM1 as their endogenous mRNAs are enriched in TGs compared to the cytosol (Supplementary Table 1). We further included MAX, known as a constitutive MYC interactor^9^, whose mRNA has no localization bias within the cytoplasm (Supplementary Table 1).

### Several MYC protein complexes are TG-dependent

Co-immunoprecipitation (co-IP) of GFP-tagged MYC in HeLa cells showed that endogenous MAX binds to MYC, regardless of whether MYC was translated in TGs or in the cytosol (Fig. 1b). In contrast, endogenous SNIP1, CycT1, or PIM1 preferentially interacted with TG-translated MYC (Fig. 1b). Next, we tested whether the presence of TGs is necessary to establish these protein complexes. We repeated the co-IP using GFP-MYC-U in cells that lack TGs through depletion of TIS11B which is required to scaffold TGs^1^. TGs were depleted using a doxycycline inducible CRISPR-Cas9 system to delete TIS11B (ED Fig. 1h). Using co-IP, we found that while the MYC-MAX interaction was TG-independent, the interactions with SNIP1, CycT1, or PIM1 were TG-dependent (Fig. 1c).

### Formation of TG-dependent MYC protein complexes requires RNA

To determine the mechanism by which TGs promote protein complex assembly, we generated recombinant proteins and reconstituted MYC protein complex assembly *in vitro* (ED Fig. 2a). We used microscale thermophoresis (MST) as a read-out to determine the binding affinities of the proteins^10^. The dissociation constant (Kd) can only be calculated if the individual measurements obtained upon titration of a binding partner can be fitted to a sigmoidal curve. As expected, GFP-tagged MAX binds MYC with a binding affinity in the low micromolar range (Fig. 1d). In sharp contrast, under the same conditions, we did not observe any measurable binding between GFP-SNIP1 and MYC (Fig. 1e, gray line).

TGs enrich a specific class of mRNAs^11^. We hypothesized that these mRNAs promote the formation of TG-dependent MYC protein complexes. To test this, we focused on the *HSPA1B* and *DNAJB1* mRNAs, which are two of the highest expressed TG-enriched mRNAs^6^. We tested whether *in vitro*-transcribed RNAs derived from the 3′UTRs of *HSPA1B* and *DNAJB1* mRNAs promote the TG-dependent interaction between GFP-SNIP1 and MYC *in vitro*. Strikingly, in the presence of either 3′UTR, MYC interacted with SNIP1 protein with micromolar affinity (Fig. 1e, pink line, ED Fig. 2b, 2c). The activity of the *HSPA1B* 3′UTR was specific for TG-dependent interactions as it did not promote the TG-independent interaction between MYC and MAX (ED Fig. 2d). In contrast, the addition of the 3′UTR of the cytosolic *SMAD6* mRNA did not promote the interaction between SNIP1 and MYC (Fig. 1f).

Surprisingly, although the *MYC* mRNA is TG-enriched, its 3′UTR did not facilitate the MYC-SNIP1 interaction (ED Fig. 2e). We previously found that TGs enrich for mRNAs that are structurally plastic and are prone to interact with other RNAs. These features can be predicted by RNAfold (Fig. ED 2f)^11^. Among the few tested mRNAs, we observed that RNAs with the highest structural plasticity score were most effective in promoting the MYC-SNIP1 protein interaction (ED Fig. 2g). Taken together, our results indicate that TG-dependent protein complexes are promoted by specific RNAs.

### mRNA-dependent protein complexes involve binding to the MYC IDR

To better understand why some MYC interactions are TG-dependent whereas others are not, we mapped the protein interaction interfaces of the four MYC interactors (Fig. 2a). This revealed that MYC interacts with MAX through the C-terminal bHLH domain, which is a folded domain^12^. In contrast, all TG-dependent protein interactors bind to the intrinsically disordered N-terminus of MYC, called here MYC IDR (Fig. 2a)^8^. The MYC IDR contains highly conserved protein binding sites, called MYC boxes I and II. They are the known binding sites for CycT1 and PIM1, whereas SNIP1 binds to the 147 N-terminal amino acids (Fig. 2a, ED Fig. 3a)^8,13^.

**Figure 2.**
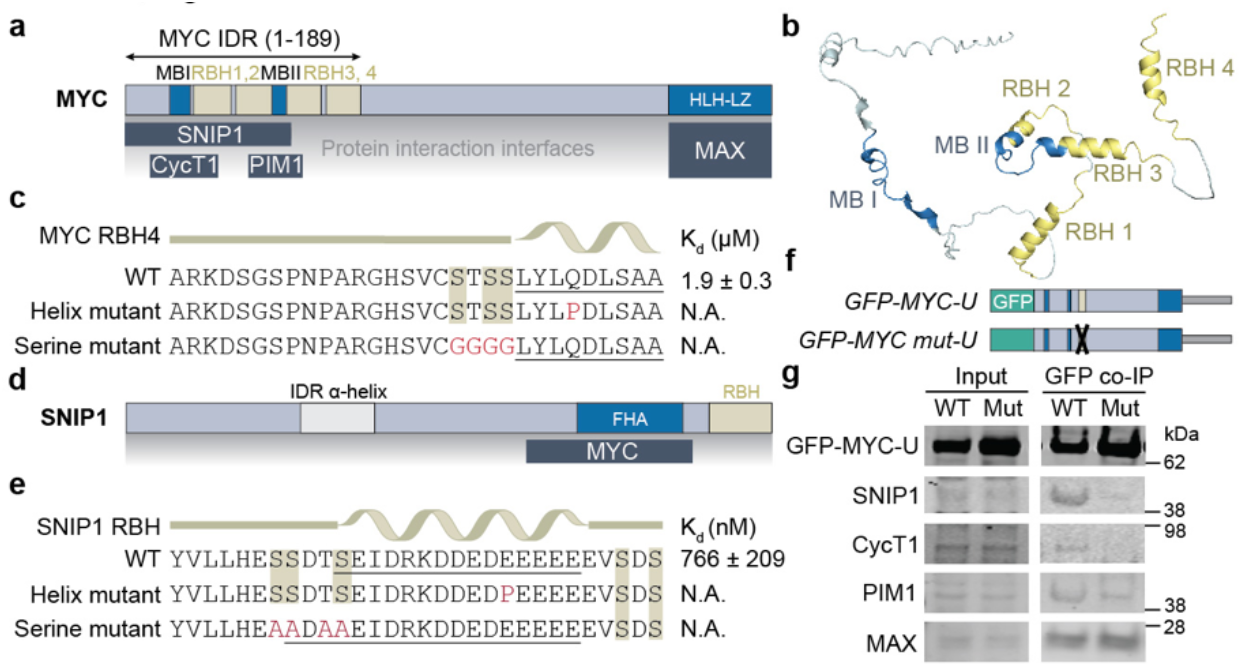
RNA-dependent protein complex assembly occurs in IDRs through a new RBD. **a**, Schematic of MYC protein domains including MYC box I (MB I), MYC box II (MB II), helix-loop-helix/leucine-zipper domain (HLH-LZ), and RNA binding helix (RBH). Shown below are the protein interaction interfaces for SNIP1, CycT1, PIM1, and MAX. **b**, AlphaFold prediction of the MYC IDR. Colors as in **a**. **c**, MST measurement of affinity of GFP-tagged MYC 30-mer peptides to the *HSPA1B* 3′UTR. Shown are peptides with wild-type (WT) sequence, helix-breaking mutation, and mutation of the flanking serines of MYC RBH4. The helix sequence is underlined. Replicates are shown in ED Fig. 3e-i. **d**, Schematic of SNIP1 protein domains. The protein interaction interface with MYC is indicated below. FHA, forkhead-associated domain. **e**, As in **c**, but the binding affinity of WT and mutant SNIP1 30-mer peptide to the *HSPA1B* 3′UTR was determined. Replicates are shown in ED Fig. 3k-m. **f**, cDNA constructs containing GFP-tagged WT and mutant MYC (RBH4 serine mutation) used for co-IP are shown. **g**, GFP co-IP of endogenous MYC interactors from HeLa cells following transfection of constructs shown in **f**.

As RNA-IDR interactions are widespread^14^, we hypothesized that the MYC IDR contains an RNA-binding domain (RBD) that allows subsequent protein complex assembly. Nearly all so far reported RNA-IDR interactions are electrostatic as they occur between positively charged amino acids and negatively charged RNAs^14-18^. However, the MYC IDR has few charged amino acids and is overall negatively charged (ED Fig. 3b)^19^. Among non-canonical RBDs that do not depend on charged amino acids, a highly conserved α-helix in the splicing factor SRSF1 was reported to bind to RNA, suggesting that α-helices may have RNA-binding capacity^20^.

### Identification of a serine-rich RNA-binding helix as new IDR RBD in transcription factors

We used AlphaFold to investigate if the MYC IDR forms transient α-helices^21-23^. AlphaFold predicts four α-helices in the vicinity of the MYC boxes I and II (Fig. 2a, 2b, ED Fig. 3c). We tested if the predicted α-helices in the MYC IDR bind to RNA. Using MST, we observed that GFP-tagged 30-mer peptides, each containing a predicted α-helix, all bind to the *HSPA1B* 3′UTR *in vitro*. Upon introduction of a helix-breaking mutation or upon mutation of the helix-flanking serine residues, RNA binding was abrogated, suggesting that both features are necessary (Fig. 2c, ED Fig. 3d-i).

MYC binds to the forkhead-associated (FHA) domain of SNIP1^13^. Adjacent to the FHA domain, SNIP1 contains a disordered C-terminus which is predicted to form an α-helix (Fig. 2d, ED Fig. 3j). Although this region is highly negatively charged, a GFP-tagged peptide containing the predicted SNIP1 α-helix bound the *HSPA1B* 3′UTR with low micromolar affinity. Again, mutation of the α-helix or the adjacent serine residues abrogated RNA binding (Fig. 2e, ED Fig. 3k-m). Taken together, both MYC and SNIP1 contain α-helices with adjacent serine residues that bind to RNA with affinities that are similar to the affinity of a classical RBD, such as the RNA-recognition motif RRM 1/2 of HuR, to its target RNA (ED Fig. 3n)^24^. In summary, we identified a new RBD in IDRs of transcription factors that we call serine-rich RNA binding helix (RBH).

Our *in vitro* binding assays suggested that TG-dependent protein complexes are RNA-dependent (Fig. 1e). Next, we tested if TG-dependent MYC protein complex assembly is RNA-dependent in cells. Three of the four RBH domains in the MYC IDR are located within the protein interaction interfaces which precludes their mutation as this may disrupt protein binding independently of RNA (Fig. 2a). We mutated RBH4 as it was located outside of the protein binding sites (Fig. 2a, 2f). We performed GFP co-IP using GFP-tagged MYC-U with a mutated RBH4 domain and observed that the mutation disrupted MYC binding to the TG-dependent interactors but had no effect on MAX binding (Fig. 2f, 2g). This result demonstrates that RNA binding to the MYC IDR is necessary to establish TG-dependent MYC protein complexes in cells.

### The MYC-SNIP1 complex forms co-translationally and acts in the nucleus

Based on our results so far, it is unclear where in cells the RNA-dependent MYC protein complexes are formed. It is commonly believed that transcription factors interact with their binding partners in the nucleus^25,26^. However, our results suggest that the MYC-SNIP1 complex assembles while MYC is translated in TGs. To investigate if the MYC-SNIP1 complex indeed forms in TGs, we performed an experiment commonly used to demonstrate co-translational protein complex assembly^27-29^. We used GFP-SNIP1 to immunoprecipitate *MYC* mRNA. If SNIP1 interacts with the MYC nascent chain during MYC translation, then GFP-SNIP1 will pull down *MYC* mRNA (Fig. 3a). To distinguish the binding of SNIP1 to *MYC* mRNA that is not associated with ribosomes, the RNA immunoprecipitation was also performed in the presence of puromycin which releases the nascent chain from the ribosome and shows the amount of *MYC* mRNA directly bound to SNIP1 (Fig. 3a)^27,28,30^. The difference between the immunoprecipitated RNA in the absence or presence of puromycin reveals the amount of SNIP1 that is cotranslationally bound. We expressed GFP-SNIP1 from a cDNA containing the *SNIP1* 3′UTR (GFP-SNIP1-U) to enable *SNIP1* mRNA localization and translation in TGs (ED Fig. 1a, 1d-f). Our results show that GFP-SNIP1 binds to MYC protein, but not to two unrelated controls, ATP5ME and RPLP0, in a co-translational manner, indicating that the MYC-SNIP1 complex forms in TGs (Fig. 3b).

**Figure 3.**
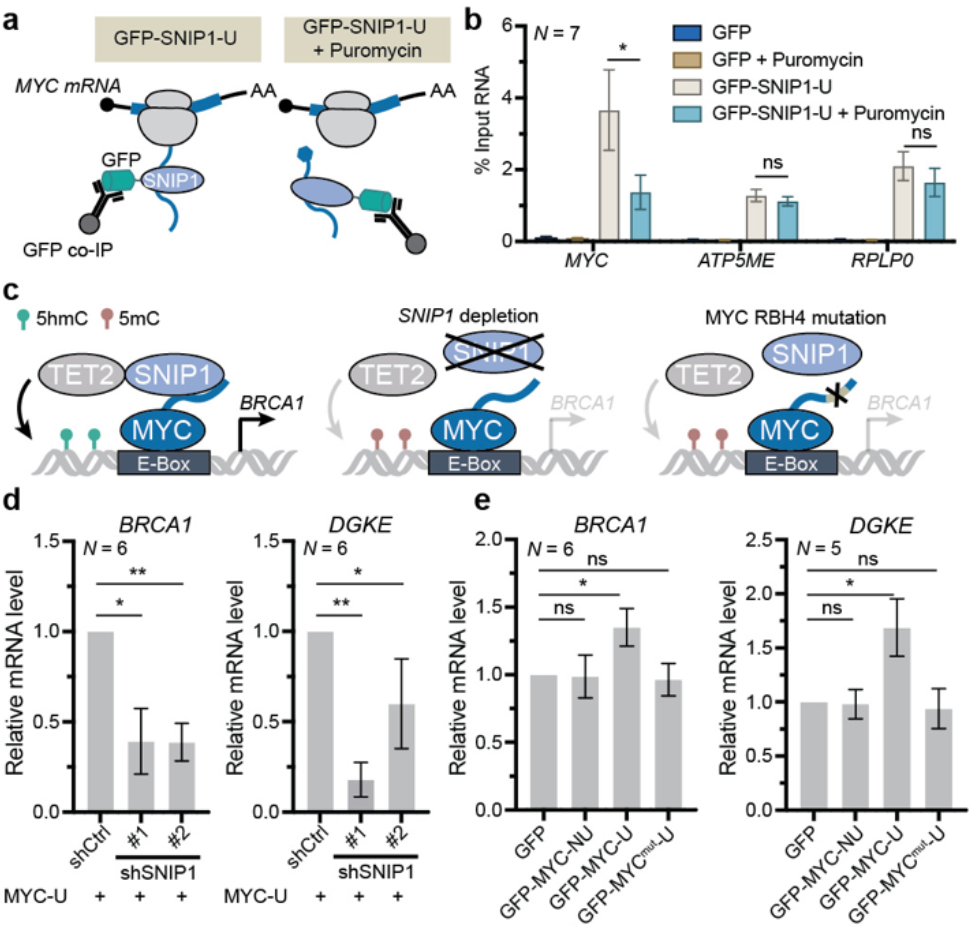
The MYC-SNIP1 complex is assembled co-translationally and acts in the nucleus. **a**, Schematic of the co-translational RNA immunoprecipitation experiment using GFP-SNIP-U. GFP-co-IP of GFP-SNIP1 in HeLa cells after transfection of a cDNA construct containing GFP-SNIP1-U. Puromycin releases the nascent chains from ribosomes, and the reduction of immunoprecipitated RNA after puromycin treatment indicates co-translational binding. **b**, Experiment as described in **a**. Further control experiments were conducted using HeLa cells transfected with GFP only, with or without puromycin treatment. *ATP5ME* and *RPLP0* are not SNIP1 targets and were used as control mRNAs. Mean ± SEM of *N = 7* biological replicates. Mann-Whitney test, *, *P* = 0.038; ns, not significant. **c**, Schematic of MYC target gene regulation by TET2-SNIP1-MYC. Left panel, loss of DNA methylation at the MYC target gene locus (*BRCA1*) upon SNIP1-dependent TET2 recruitment promotes MYC-dependent *BRCA1* expression. Middle and right panels, *SNIP1* depletion or MYC RBH4 mutation disrupts SNIP1-dependent TET2 recruitment, resulting in decreased expression of MYC target genes. 5hmC, 5-hydroxymethylcytosine; 5mC, 5-methylcytosine. **d**, mRNA expression of MYC target genes *BRCA1* and *DGKE* normalized to *ACTB* was obtained by qRT-PCR. GFP-MYC-U is transfected (abbreviated as MYC-U). Shown is mean ± SEM of six biological replicates obtained from U2OS cells. Mann-Whitney test, *, *P* = 0.049; **, *P* = 0.003. **e**, mRNA expression of MYC target genes *BRCA1* and *DGKE* normalized to *ACTB* was obtained by qRT-PCR. Shown is mean ± SEM of biological replicates obtained from U2OS cells after transfection of the indicated cDNA constructs. GFP-MYC-mut-U is RBH4 serine mutant. Unpaired t-test, *, *P* = 0.03; ns, not significant.

MYC is a transcription factor and performs its functions in the nucleus^7,8^, therefore, we examined if the TG-dependent MYC-SNIP1 complex acts in the nucleus. To do so, we investigated if the MYC-SNIP1 complex induces MYC target genes whose expression is not increased by MYC-NU, which is translated in the cytosol and is unable to bind to SNIP1. TET2 is a DNA dioxygenase that catalyzes demethylation of 5-methyl cytosine at enhancers to promote transcriptional activity through loss of DNA methylation^31^. TET2 lacks a DNA binding domain and uses adaptors for recruitment to specific genomic loci. It was shown previously that SNIP1 acts as adaptor that recruits TET2 to MYC to control MYC target gene expression (Fig. 3c)^13,32^.

We confirmed that in cells expressing MYC-U, knockdown of SNIP1 decreases the expression of the MYC target genes *BRCA1* and *DGKE* (Fig. 3d, ED Fig. 4a)^32^. A similar phenotype was obtained upon disruption of MYC-SNIP1 complex assembly by mutating MYC RBH4 which prevents MYC binding to RNA. The RBH4 mutant decreased MYC target gene expression compared with the wild-type MYC-U (Fig. 3c, 3e). Importantly, lower MYC target gene expression was also observed with MYC-NU which is translated in the cytosol and cannot bind to SNIP1 (Fig. 3e). This experiment demonstrates that the MYC-SNIP1 complex formed in TGs and is functional. It also suggests that MYC-NU cannot bind to SNIP1 in the nucleus, indicating that the RNA-dependent MYC-SNIP1 complex can only form cotranslationally and not in a post-translational manner in cells. Our data show that translation of MYC in two different cytoplasmic compartments results in at least two different MYC protein states that are distinguished by the presence or absence of the RNA-dependent MYC interactor SNIP1.

**Figure 4.**
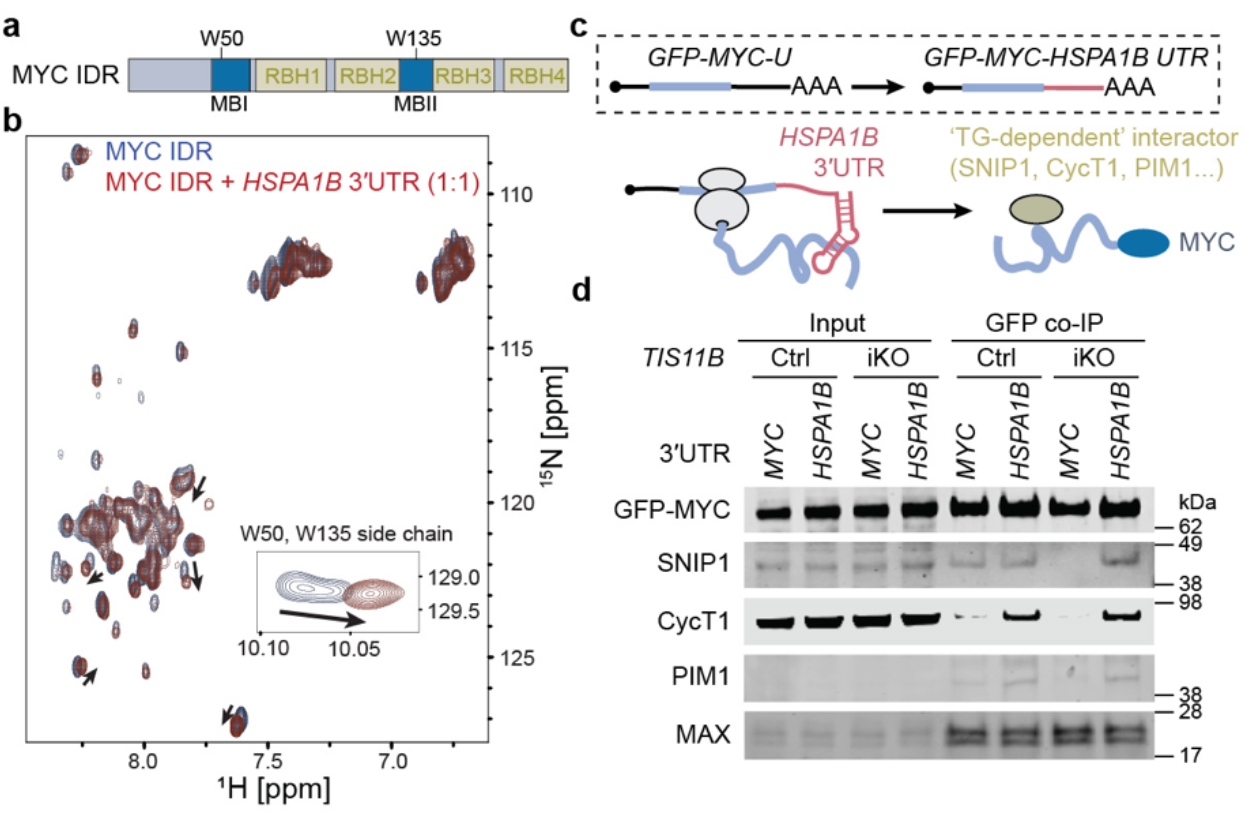
RNA binding to the MYC IDR changes the IDR conformational ensemble and is sufficient for MYC protein complex assembly in cells. **a**, Schematic of the MYC IDR used for NMR. Indicated are the two tryptophan residues (W50, W150) of the MYC IDR that are located in the protein binding interfaces. **b**, ^1^H^15^N HSQC spectra of the MYC IDR in the absence (blue) or presence of RNA (red, *HSPA1B* 3′UTR) at 1:1 protein-RNA ratio. **c**, cDNA constructs to test if a 3′UTR with IDR chaperone activity is sufficient for RNA-dependent MYC protein complex assembly in the absence of TGs. Shown is the *MYC* mRNA that serves as control and a chimeric mRNA containing the MYC coding sequence with the *HSPA1B* 3′UTR as a 3′UTR with IDR chaperone activity. A model for IDR chaperone activity *in cis* based on the chimeric cDNA construct is described below. **d**, GFP co-IP of endogenous MYC interactors after transfection of the cDNA constructs from **c** in control or *TIS11B* iKO HeLa cells.

### RNA interaction changes the IDR conformation in the region of the protein binding sites

Next, we addressed the molecular mechanism by which RNA enables the MYC IDR to engage in protein complex assembly. Previous structural analyses revealed that, in most cases, RNA binding to disordered protein regions results in a conformational change of the RNA as well as the RNA-binding protein^24,33-37^. We used NMR spectroscopy to test the hypothesis that RNA interaction changes the conformational ensemble of the MYC IDR. For these studies, we used a construct containing the IDR of MYC (Fig. 4a, ED Fig. 2a).

Indeed, addition of the *HSPA1B* 3′UTR resulted in small but significant perturbations in the 2D ^1^H-^15^N HSQC spectra of the ^15^N labeled MYC IDR (Fig. 4b, arrows). Interestingly, the largest perturbation was observed for the tryptophan side chains (Fig. 4b, insert). Intriguingly, the two tryptophans of the MYC IDR are located in MYC boxes I and II which are the protein binding sites (Fig. 4a). These results strongly suggest that RNA binds to the MYC IDR region and likely changes the conformational properties of regions involved in protein-protein interaction.

### RNA binding is necessary and sufficient for formation of MYC IDR protein complexes

Based on our results, we propose that the *HSPA1B* 3′UTR has IDR chaperone activity, as it binds to the MYC IDR, changes the IDR conformational ensemble, and allows MYC protein complex assembly (Fig. 2c, 2g, 4b). These results indicate that RNA binding is necessary for TG-dependent protein complex assembly.

To examine if RNA binding is sufficient for TG-dependent protein complex assembly in cells, we expressed MYC from a cDNA containing the *MYC* coding region and replaced the *MYC* 3′UTR with the *HSPA1B* 3′UTR, which has IDR chaperone activity (Fig. 4c). Co-IP of MYC expressed from the chimeric mRNA showed that the presence of the *HSPA1B* 3′UTR was sufficient to induce protein complex assembly of MYC-SNIP1, MYC-CycT1, and MYC-PIM1 (Fig. 4d). Importantly, the *HSPA1B* 3′UTR promoted MYC protein complex assembly even in the absence of TGs (Fig. 4d). This result demonstrates that binding of RNA with IDR chaperone activity is necessary and sufficient for TG-dependent protein complex assembly involving the MYC IDR.

### The serine-rich RNA binding helix is a widespread RBD in IDRs

Next, we asked if RNA-dependent protein complex assembly is used by transcription factors other than MYC. As RNA-dependent protein complex assembly occurs in IDR regions, we predicted IDRs and examined if their occurrence is overrepresented in TG- or cytosolically-translated proteins. We observed that proteins encoded by mRNAs with a biased localization to TGs contain more and larger IDRs than cytosolically-translated proteins (Fig. 5a, ED Fig. 5a). These results suggest that protein complex assembly involving IDRs is widespread in TGs.

**Figure 5.**
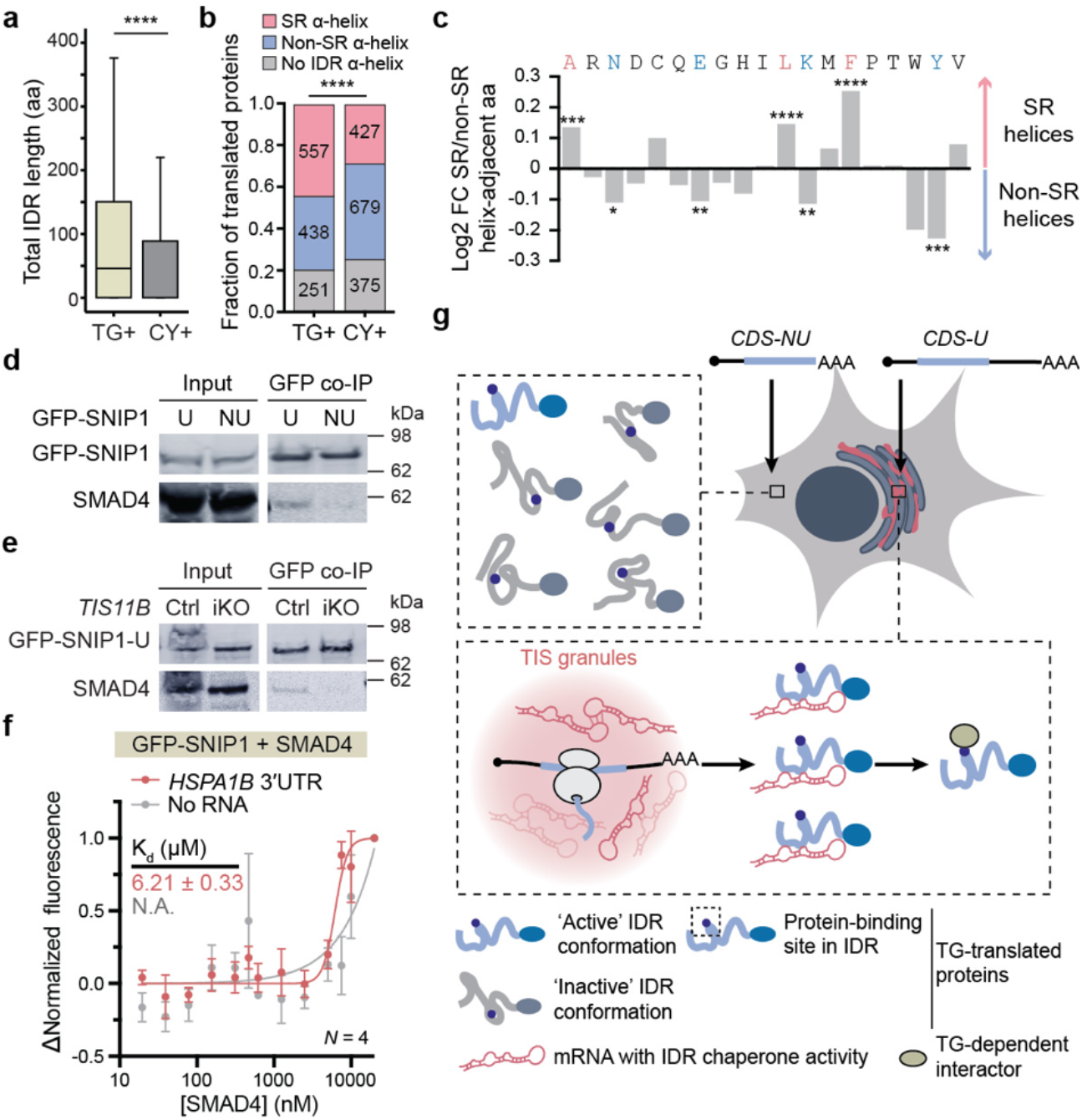
RNA-mediated protein complex assembly involving serine-rich RBH domains in IDRs is widespread. **a**, The total amino acid (aa) length of IDRs is shown for proteins encoded by mRNAs enriched in TGs (TG+, *N =* 1246) or the cytosol (CY+, *N =* 1481)^6^. Only IDRs with ≥ 30 amino acids were included. Mann-Whitney test, ****, *P* = 2.1E-21. **b**, As in **a**, but the fraction of proteins translated in a compartment-biased manner that contain different types of IDR α-helices are shown. Chi-square test (χ^2^ = 62.8) was performed to test enrichment of serine-rich (SR) α-helices over non-serine-rich (non-SR) α-helices between the indicated groups. ****, *P* < 0.0001. **c**, Amino acid enrichment in the helix-flanking region (5 aa up- or downstream) in serine-rich (SR) RBH domains (*N* = 6345) compared with IDR α-helices without enrichment of serines in the helix adjacent regions (non-SR, *N* = 42039). Chi-square test was performed. A, ***, *P* = 9E-4; N, *, *P* = 0.02; E, **, *P* = 0.008; L, ****, *P* < 0.0001; K, **, *P* = 0.004; F, ****, *P* < 0.0001; Y, ***, *P* = 8E-4. Amino acids that are not significantly enriched in either group are not labeled. **d**, GFP co-IP of endogenous SMAD4 after transfection of cDNA constructs containing GFP-SNIP1-U or GFP-SNIP1-NU in HeLa cells. **e**, GFP co-IP of endogenous SMAD4 after transfection of cDNA constructs containing GFP-SNIP1-U in control or *TIS11B* iKO HeLa cells. **f**, Mean ± std of MST measurement of the SNIP1-SMAD4 interaction *in vitro* in the absence or presence of 200 nM *HSPA1B* 3′UTR. **g**, Model for TG-dependent protein complex formation. 3′UTR-dependent mRNA localization to TGs. TG-enriched mRNAs bind to the newly translated IDR and promote an active IDR conformation. When translated in the cytosol, the IDR is predominantly present in an inactive conformational state which prevents interactor binding.

Next, we set out to identify additional TG-translated proteins with RBH domains. As the α-helices of the MYC and SNIP1 RBH domains were predicted by AlphaFold^22,23^, we identified all AlphaFold-predicted α-helices located outside of known folded domains according to UniProt. Among the TG-translated proteins CycT1, PIM1, RBL1, and SMAD4, we observed several predicted IDR α-helices and tested if they bind to the *HSPA1B* 3′UTR (ED Fig. 5b)^6^. Using MST, we measured the affinity of RNA to GFP-tagged 30-mer peptides containing these α-helices (ED Fig. 5b-m). Among 13 tested α-helices, all (*N* = 7) that contained at least two serines in the five amino acids flanking the predicted α-helix bound to the *HSPA1B* 3′UTR, whereas all other α-helices did not bind (ED Fig. 5c, 5d).

These results suggest that serine-rich RBH domains bind to TG-enriched RNAs, such as the *HSPA1B* 3′UTR. Overall, our predictions identified 16319 serine-rich RBH domains present in 7921 proteins in the UniProt proteome, suggesting that this new RBD is widespread (Supplementary Table 2). Moreover, the serine-rich RBH domains are significantly enriched among TG-translated proteins compared with predominantly cytosolically-translated proteins (Fig. 5b).

The RNA binding capability of non-serine-rich α-helices is currently unknown. However, these helices contain a significant enrichment of lysines and tyrosines within their flanking regions (Fig. 5c). This is consistent with a previously observed overrepresentation of these amino acids in RNA-binding IDRs^14^. Therefore, it is likely that a sizeable fraction of non-serine-rich IDR α-helices will bind to RNA, but the features of the RNA targets are currently unknown.

Lastly, we investigated if transcription factors other than MYC use mRNA binding in TGs for protein complex assembly. SNIP1 is known to form a complex with SMAD4, and both *SNIP1 and SMAD4* mRNA transcripts are enriched in TGs^6,38^. GFP co-IP showed that formation of the SNIP1-SMAD4 complex in cells required the presence of the *SNIP1* 3′UTR (Fig. 5d). As SNIP1-NU is translated in the cytosol and SNIP1-U is predominantly translated in TG (ED Fig. 1a, 1d-f), these results suggest that TG-translated SNIP1 interacts with endogenous SMAD4 (Fig. 5d). Moreover, the SNIP1-SMAD4 interaction was TG-dependent in cells and RNA-dependent *in vitro*, demonstrating that additional transcription factors use RNA-dependent protein complex assembly in TGs (Fig. 5e, 5f, ED Fig. 5n).

### Model of mRNA-dependent assembly of protein complexes involving IDRs

Taken together, we propose the following model for the formation of mRNA-dependent protein complexes. mRNAs use TIS11B binding sites in their 3′UTRs to localize to TGs. During translation of proteins with IDRs, TG-enriched mRNAs with IDR chaperone activity are in close proximity when the IDR in the nascent chain becomes exposed from the ribosome. These mRNAs bind to the IDR, change the conformational ensemble of the IDR, and drive subsequent protein complex assembly (Fig. 5g). Our current data suggest that TG-translated proteins mostly use RNA binding by the surrounding TG-enriched mRNAs *in trans* to induce a conformational change, as, for example, the *MYC* 3′UTR did not have IDR chaperone activity for the MYC IDR (ED Fig. 2e). Nevertheless, 3′UTRs with IDR chaperone activity can act *in cis*, thereby overcoming the requirement for the presence of TGs (Fig. 4d). These results suggest that the essential function of TGs in the regulation of protein complex assembly is to provide a high concentration of mRNAs with IDR chaperone activity that promote active IDR conformations during translation.

## Discussion

### mRNAs regulate activity of proteins with IDRs

Protein activity is primarily regulated by ligand binding and post-translational modifications (PTMs). While ligands mostly affect activity of folded proteins^39,40^, PTMs are often critical to switch on proteins with IDRs^41,42^. For example, proteins whose binding sites are not accessible by default are autoinhibited. Autoinhibition is most often overcome through the addition of PTMs leading to conformational changes that expose active sites or ligand binding sites^41^. Here, we found that RNA binding controls protein-protein interactions and the activity of proteins with IDRs. IDRs are widespread, as 63% of human proteins contain disordered regions^43^. They are particularly enriched in regulatory factors, such as transcription factors and enzymes^44^. Moreover, RNA-IDR interactions are widespread as half of all RNA-protein interaction events in cells are not accomplished by canonical RBDs, but by IDRs^14^. These findings imply that RNA has the potential to be a major regulator of protein function.

### mRNAs regulate protein activity by acting as IDR chaperones

In addition to the known function of mRNAs as templates for protein synthesis, we reveaIed here a new roIe: We propose that certain mRNAs have IDR chaperone activity as they bind to IDRs and change their conformational ensembles, thereby promoting subsequent protein complex assembly. The exact biophysical features that allow mRNAs to act as IDR chaperones are currently not known. Our data suggest that these mRNAs are enriched in TGs. Moreover, we observed that RNAs with a high score for structural plasticity have more capacity to induce changes in the IDR conformational ensemble (ED Fig. 2g), suggesting that structural plasticity may be important for IDR chaperone activity.

mRNAs with a high structural plasticity score were previously identified, as they are responsible for the characteristic network-like morphology of TGs. It was shown that mRNAs with a high structural plasticity score form extensive RNA-RNA interactions^11^. Our data suggest that this feature may also promote RNA interaction with IDRs. With respect to the molecular mechanism of chaperoning IDRs, it is possible that RNAs may change the solvent environment of the IDRs, may influence the dynamics of conformational transitions, or may induce disorder to order transitions, thus exposing the protein binding sites^42,44-49^. Identifying the molecular mechanism of action of IDR chaperones is important as it will facilitate the development of biotechnology applications that use RNA to control protein activity.

### Translation in TGs enables different protein functional states

Our current model of TG-dependent protein complex assembly involves the following steps. mRNA localization to different subcytoplasmic compartments is influenced by a combinatorial code of 3′UTR-bound RNA-binding proteins^6^. mRNAs use TIS11B binding sites in their 3′UTRs to localize to TGs, where they are translated^1,6^. The surrounding TG-enriched mRNAs bind to newly translated IDRs inducing a change in the conformational ensemble which drives subsequent protein complex assembly. Our current data suggest that TGs provide a high concentration of mRNAs with IDR chaperone activity. Although these mRNAs are present in the cytosol, they do not achieve sufficient proximity or concentration to affect IDR conformational ensembles.

Most mRNAs are translated in at least two subcytoplasmic environments^6^. Our data imply that translation in different compartments allows MYC protein to be present in at least two different protein states, despite having an identical amino acid sequence. Cytosolically-translated MYC is present in MYC protein state 1, which binds to MAX but favors an inactive IDR conformation.

MYC protein state 1 is not able to bind to SNIP1 and cannot recruit TET2 to promote MYC target gene expression. In contrast, TG-translated MYC is present in MYC protein state 2, which binds MAX and favors an active IDR conformation, allowing it to bind to SNIP1 and induce specific MYC target genes. Our data suggest that 3′UTR-dependent translation in different subcytoplasmic compartments allows a protein to have different functions within one cell type, despite using the same protein sequence.

## Acknowledgements

This work was funded by the NIH Director′s Pioneer Award (DP1-GM123454), an R35GM144046 NIH grant, a grant from The G. Harold and Leila Y. Mathers Foundation, a grant from The Pershing Square Foundation, William Ackman, and Neri Oxman, and the MSK Core Grant (P30 CA008748) to CM. H.M.A-H. received funding from NIH (NIGMS: R01GM132899). H.M.A-H is a member of the New York Structural Biology Center. The NMR data collected at NYSBC was made possible by an ORIP/NIH facility improvement grant CO6RR015495. We thank Xiuzhen Chen for critical reading of the manuscript and all members of the Mayr Lab for helpful discussions.

## Author contributions

Y.L. performed all experiments, except the RNA-FISH assays which were performed by E.H.L. S.P. and H.M.A-H performed the NMR experiments. A.S.J.S.M. computationally identified the IDR helices. S.M. generated the Cas9 expressing cell lines. Y.L. and C.M. conceived the project, designed the experiments, and wrote the manuscript with input from all authors.

## Methods

### Cell lines

The HeLa cell line was a gift from the lab of Jonathan S. Weissman (UCSF). The U2OS cell line was a gift from the lab of Thijn Brummelkamp (Netherlands Cancer Institute). Both cell lines were maintained at 37°C with 5% CO2 in DMEM containing 4500 mg/l glucose, 10% heat-inactivated fetal bovine serum, 100 U/ml penicillin, and 100 mg/ml streptomycin. The cell lines have not been authenticated.

### Constructs

Unless otherwise stated, all mammalian expression vectors were derived from pcDNA3.1-puro mGFP (monomeric GFP, A207K), which was reported previously^1^. All coding sequences were PCR-amplified from HeLa cDNA, and 3′UTR sequences were from HeLa genomic DNA. All primer sequences are listed in Supplementary Table 3. All constructs were sequence verified.

#### GFP-MYC

To generate GFP-MYC-NU, the MYC coding sequence (1365 bp) was PCR-amplified and inserted between *BsrGI* and *BamHI* sites. To generate GFP-MYC-U, the *MYC* 3′UTR (473 bp) was amplified and inserted between *BamHI* and *EcoRI*. To obtain the GFP-MYC-mut-U construct, the MYC serine mutation insert was generated by overlap extension PCR where the RBH4 mutation (STSS to GGGG or STSS to AAAA) was incorporated into the reverse or forward primers of each piece and then subcloned between *BsrGI* and *BamHI* sites. To generate the chimeric construct containing the MYC coding region followed by the *HSPA1B* 3′UTR (GFP-MYC-HSPA1B-UTR), the *HSPA1B* 3′UTR (379 bp) was inserted downstream of the MYC coding region in GFP-MYC-NU and the BGH polyA site (nucleotides 1025-1252 in pcDNA3.1 puro) was deleted by inverse PCR.

#### GFP-SNIP1

To prepare the GFP-SNIP1-NU construct, the SNIP1 coding sequence (1188 bp) was cloned between *BsrGI* and *XhoI* using compatible cohesive ends generated by *BsiWI*. To obtain the GFP-SNIP1-U construct, the short 3′UTR (1212 bp) was inserted between *XbaI* and *ApaI*. Then, the polyadenylation site of the short 3′UTR was mutated to AAGCAA, and the remaining sequence from the long 3′UTR (2087 bp) was inserted between *ApaI* and *PmeI*.

#### SNIP1 knockdown

For shRNAs that knockdown SNIP1, pLKO.1 puro was used. The forward sequences of the synthetic DNA oligonucleotides are listed in Supplementary Table 3 and were inserted into pLKO.1 puro between *SgrAI* and *EcoRI*. The shRNAs used were either purchased from Addgene (shCtrl: Addgene #1864) or designed using the Genetic Perturbation Platform.

#### Recombinant proteins

The pET28a vector containing a 6×His-MBP tag was previously reported and was used to clone all constructs for recombinant protein expression^11^. The monomeric GFP tag was inserted between *NheI* and *BamHI* to prepare all N-terminal GFP fusion proteins used in MST experiments. In general, GFP fusions of full-length proteins contain a C-terminal Strep-Tag II (SAWSHPQFEK), but those of 30-mer peptides do not. The Strep-Tag II, where indicated, was included in the reverse primer used to amplify the inserts.

6×His-MBP-GFP-MAX-Strep-Tag II was obtained by cloning the MAX coding sequence (480 bp) between *BsrGI* and *BamHI*. 6×His-MBP-GFP-SNIP1-Strep-Tag II was obtained by inserting the SNIP1 coding sequence (1188 bp) between *BsrGI* and *XhoI* using compatible cohesive ends generated by *BsiWI*. 6×His-MBP-GFP-HuR RRM1/2-Strep-Tag II was prepared by inserting the GFP-HuR RRM1/2 (HuR amino acids 19-189) between *NheI* and *BamHI*. 6×His-MYC-Strep-Tag II was obtained by inserting the MYC coding sequence (1317 bp) between *NheI* and *BamHI* sites. 6×His-MYC IDR was cloned by PCR-amplifying the 2-189 amino acids of MYC isoform 1 (564 bp) and inserting it between *NheI* and *BamHI*. 6×His-SMAD4 was produced by cloning the SMAD4 coding sequence (1656 bp) between *NheI* and *HindIII*. All primers are listed in Supplementary Table 3.

6×His-MBP-GFP fusions of 30-mer peptides were created by inserting sequences encoding each region between the *BsrGI* and *BamHI* sites, unless otherwise indicated. MYC RBH4 helix mutation and serine mutation sequences were amplified from the corresponding pcDNA3.1-GFP-MYC mutant constructs. 6×His-MBP-GFP-SNIP1 RBH was generated by inserting SNIP1 RBH 30-mer between *BsrGI* and *XhoI*. RBH mutant inserts of SNIP1 were obtained as synthetic oligos from Genewiz and cloned into the pET28a-6×His-MBP-GFP construct using the same sites. All 30-mer peptide sequences that were fused to 6×His-MBP-GFP are listed in Supplementary Table 3.

#### DNA templates for RNA *in vitro* transcription

All DNA templates were PCR-amplified and purified by gel extraction. The DNA sequence of the 3′UTR of *MYC* (473 bp) was amplified from the pcDNA3.1-GFP-MYC-U construct. The 3′UTRs of *HSPA1B* (379 bp), *DNAJB1* (1174 bp), and *SMAD6* (469 bp) were PCR-amplified from HeLa genomic DNA. The T7 promoter (TAATACGACTCACTATAGGG) was incorporated into the forward primers used for generating these templates. All primers used for preparing the DNA templates are listed in Supplementary Table 3.

### Antibodies

Primary antibodies used in this study include the following: chicken anti-GFP (ab13970, Abcam), rabbit anti-ZFP36L1/2 (2119, Cell Signaling Technology), rabbit anti-SNIP1 (14950-1-AP, Proteintech), mouse anti-GAPDH (G8795, Sigma Aldrich), rabbit anti-SMAD4 (ab40759, Abcam), rabbit anti-CCNT1 (GTX133413, Genetex), mouse anti-MYC (sc-42, SCBT), mouse anti-PIM1 (sc-13513, SCBT), goat anti-MAX (AF4304, R&D systems).

The following secondary antibodies were used in this study: donkey anti-chicken IgG IRDye 680 (926-68075, LI-COR), goat anti-mouse IgG IRDye 680 (926-68070, LICOR), goat anti-rabbit IgG IRDye 800 (926-32211, LI-COR), donkey anti-goat IgG IRDye 800 (926-32214, LI-COR).

### RNA-FISH

RNA-FISH experiments probing for GFP-fusion constructs were performed as described previously^1^. Stellaris FISH probes for eGFP with Quasar 670 Dye were used. HeLa cells were seeded on 4-well Millicell EZ slides (Millipore), and GFP fusion constructs encoding cDNAs of interest were transfected into HeLa cells using Lipofectamine 3000. 100 ng BFP-TIS11B was co-transfected to label TIS granules. 24 h after transfection, cells were washed with PBS for 5 min, fixed with 4% paraformaldehyde for 10 min at room temperature, and washed twice with PBS. PBS was aspirated, and cells were permeabilized with 1 ml 70% ethanol at 4 °C for 2-8 hours. The 70% ethanol was then discarded, and 1 ml wash buffer (2×SSC, 10% formamide in nuclease-free water) was added to each well and incubated for 5 min at room temperature.

Hybridization mix was prepared by mixing 10% Dextran sulfate, 10% formamide, 2×SSC, 2 mM ribonucleoside vanadyl complex, 0.02% BSA, 200 μg/ml yeast tRNA, 200 μg/ml single strand DNA, and FISH probe against GFP (1:200). 200 μl hybridization mix was added to each well and incubated at 37 °C overnight. Slides were washed twice with 1 ml pre-warmed wash buffer at 37 °C in the dark, 30 min each. Slides were then washed with PBST and mounted with ProLong Gold Antifade mounting solution (Invitrogen). Images were captured using confocal ZEISS LSM 880 with Airyscan super-resolution mode.

#### Line profile analysis

Line profiles were generated with FIJI (ImageJ). Two to four straight lines were drawn across TIS granules in different directions for each cell, indicated by arrows in each figure. Fluorescence intensity along the straight line of BFP-TIS11B protein and co-transfected target RNAs were calculated using the FIJI plot profile tool. The Pearson’s correlation coefficient (R) of two fluorescent signals was calculated with Excel. R = 1 indicates perfect colocalization of the mRNA and the TIS granules, while R = -1 indicates complete exclusion of the mRNA from TIS granules. R = 0 suggests a random distribution of the mRNA across the cytosol and TIS granules.

### Generation of doxycycline inducible *TIS11B* knockout cell line (*TIS11B* iKO)

Doxycycline inducible Cas9 (iCas9) HeLa cells were generated by infecting cells with lentivirus containing a Cas9-P2A-GFP expression cassette under a doxycycline inducible promoter as described previously (Addgene plasmid #85400)^50^. During consecutive rounds of fluorescence-activated cell sorting, we selected a cell pool exhibiting robust induction of Cas9/GFP expression after doxycycline treatment (100 ng/ml for 24 hours), and low levels of leaky transgene expression in the absence of the drug.

Next, we transduced iCas9 cell lines with a lentiviral construct harboring a pair of guide RNAs either targeting *TIS11B* or non-targeting control gRNAs. To generate these constructs, we adapted the plentiGuide-puro vector (Addgene plasmid # 52963)^51^ to incorporate a second guide RNA expression cassette as described previously^52^. For this purpose, the plasmid was digested with BsmBI (FastDigest Esp3I, Thermo Fisher Scientific) and a synthetic 391 bp double-stranded DNA fragment encoding 5′-(1st gRNA/scaffold/H1 promoter/2nd gRNA)-3′ was inserted using the NEBuilder HiFi assembly system (NEB). Synthetic DNA fragments were ordered from Genewiz and sequences are listed in Supplementary Table 3. The assembled vector DNA was used to transform chemically competent Stbl3 bacteria cells (Invitrogen), and correct vector clones were identified by Sanger sequencing.

Lentivirus was generated in HEK293T cells using standard methods and 200 μl of viral supernatant was used to transduce iCas9 cells in a 6-well dish together with 8 μg/ml polybrene. Transduced cells were subjected to puromycin selection (1 μg/ml) for five days and resistant cells were aliquoted and frozen for all further experiments. Finally, for induction of gene knockouts, *TIS11B* iKO and corresponding control cells (containing non-targeting control guide RNAs) were treated with doxycycline (100 ng/ml) for up to seven days, after which TIS11B protein expression was evaluated by western blotting.

### Co-immunoprecipitation (co-IP)

mGFP-MYC or mGFP-SNIP1 fusion constructs were transfected into HeLa cells and co-IP of endogenous protein interactors was performed. Alternatively, transfection was performed using HeLa/iCas9 cells treated with 100 ng/ml doxycycline for 6 days. Cells were lysed 24 h after transfection using 250 μl RIPA buffer (10 mM Tris-Cl pH 7.5, 150 mM NaCl, 0.5 mM EDTA, 0.1% SDS, 1% Triton X-100, 1% deoxycholate) supplemented with protease inhibitor cocktail (Roche). Cell lysates were incubated on ice for 30 min, followed by 30 s sonication on ice with on/off intervals of 1 and 2 s. Cell lysates were then spun down at 20,000 g in a microcentrifuge for 10 min at 4 °C. The supernatant was transferred to a pre-chilled Eppendorf tube and diluted with 350 μl GFP-Trap dilution buffer (10 mM Tris-Cl pH 7.5, 150 mM NaCl, 0.5 mM EDTA). 20 μl of the diluted lysate was mixed with 4× Laemmli sample buffer (0.2 M Tris-Cl, 0.4 M DTT, 8% (w/v) SDS, 4.3 M glycerol, 6 mM bromophenol blue) and saved as the input sample. The remaining diluted lysate was then mixed with 10 μl GFP-Trap agarose beads (Chromotek) that were pre-equilibrated with GFP-Trap dilution buffer. The lysate-bead mix was rotated at 4 °C for 2 h. The GFP-Trap beads were spun down at 2,500 g for 5 min at 4 °C and washed three times with GFP-Trap dilution buffer. A 2× Laemmli sample buffer was added to the beads, boiled at 95 °C for 7 min, spun down at 2,500 g for 2 min, and cooled on ice before loading on a NuPAGE 4-12% Bis-Tris gradient protein gel (Invitrogen). Western blotting was performed following the manufacturer’s instructions, and the image was captured using the Odyssey DLx system (LI-COR).

### *In vitro* transcription of RNA

All RNAs were prepared by *in vitro* transcription using the T7 MEGAscript kit (Invitrogen). 500 ng of PCR-amplified DNA template was used for each 20 μl reaction. The reaction mix was prepared following the manufacturer’s protocol and incubated at 37 °C for 6 h. The products were then treated with Turbo DNase provided in the kit and incubated at 37 °C for 30 min. The integrity and sizes of the RNAs were examined using agarose gels. All RNAs were precipitated with LiCl overnight at -20 °C. The RNAs were centrifuged at 20,000 g for 30 min at 4 °C. The pellets were washed with 1 ml 70% ethanol, redissolved in nuclease-free water, aliquoted, and stored at -20 °C. The concentration of RNA was measured by NanoDrop.

### Recombinant protein expression and purification

All expression constructs were transformed into *BL21* E. coli (NEB) for the following protein expression steps.

#### 6×His-MBP-GFP and Strep-tag II dual-tagged proteins

To purify 6×His-MBP-GFP and Strep-tag II dual-tagged proteins, we used two steps of purification. Specifically, a fresh colony was used to inoculate an overnight culture in 5 ml LB/kanamycin media and shaken vigorously at 250 rpm at 37 °C. The overnight culture was diluted in 500 ml LB media containing 50 mg/l kanamycin and grown at 37 °C until OD600 reached 0.6-0.8. Protein expression was induced by adding 0.5 mM IPTG, and the bacteria were cultured at 18 °C overnight.

Bacteria were harvested by centrifugation at 6,000 g for 15 min. The pellet was resuspended in 30 ml cold lysis buffer (50 mM sodium phosphate pH 7.4, 600 mM NaCl, 1 mM PMSF), supplemented with 10 mg lysozyme (Life Technologies), one tablet of protease inhibitor cocktail (Roche) and 1 mg DNase I (Roche). After incubation on ice for 30 min, bacteria were sonicated on ice for 3 min 30 seconds with on/off intervals of 1 and 3 s. The lysate was cleared by centrifugation at 38,000 g at 4 °C for 30 min.

5 ml TALON metal affinity resin (TaKaRa Bio) was equilibrated with five column volumes of native lysis buffer. The supernatant of cleared bacteria lysate was transferred to a new 50 ml Falcon tube and incubated with TALON resin with rotation at 4 °C for 2 h. The slurry was transferred into a gravity column and washed with 10 column volumes of native wash buffer (50 mM sodium phosphate pH 7.4, 600 mM NaCl, 10 mM imidazole). The protein was eluted with five column volumes of elution buffer (50 mM sodium phosphate pH 7.4, 600 mM NaCl, 200 mM imidazole).

The eluted protein was mixed with 1 ml Strep-Tactin Superflow agarose (Qiagen) and incubated at 4 °C for 30 min. The mixture was transferred to a gravity column, washed with 5 column volumes of wash buffer 2 (20 mM Tris-Cl pH 7.4, 600 mM NaCl), and eluted with 5 column volumes of elution buffer 2 (20 mM Tris-Cl pH 7.4, 600 mM NaCl, 10 mM desthiobiotin). The integrity of the purified protein was examined by SDS-PAGE and Coomassie blue staining. The oligomeric state of the protein was evaluated by 4-16% Bis-Tris gradient NativePAGE gel (Invitrogen) and Coomassie staining.

The following secondary antibodies concentrated, and the buffer was exchanged using centrifugal filters with a 30 kDa cutoff (Pall). Protein was aliquoted and stored in 25 mM Tris-Cl pH 7.4, 150 mM NaCl and 0.01% Tween-20 at -80 °C.

#### 6×His-MBP-GFP-tagged 30-mers

6×His-MBP-GFP-tagged 30-mers were purified using TALON resin under native conditions which uses the first step of purifying the dual-tagged constructs.

##### SMAD4

6×His-tagged full-length SMAD4 was purified using TALON resin under native conditions.

##### MYC and MYC IDR

6×His-tagged full-length MYC and MYC IDR (2-189) were purified using a modified protocol from previous reports^53^. Briefly, 1 liter of bacteria culture was induced with 0.5 mM IPTG when OD600 reached 0.4 and shaken at 30 °C for 3 h before harvest. The cell pellet was resuspended in 30 ml ice-cold lysis buffer 2 (20 mM HEPES pH 7.4, 500 mM NaCl, 10% (v/v) glycerol, 0.1% (v/v) NP-40, 1 mM PMSF) supplemented with lysozyme and protease inhibitors. The suspension was incubated on ice and sonicated for 3 min 30 seconds with on/off intervals of 1 and 3 s. The lysate was centrifuged at 38,000 g at 4 °C for 30 min. The supernatant was decanted, and the cell pellet was extracted with 10 ml buffer E (50 mM HEPES pH 7.4, 5% (v/v) glycerol, 1% (v/v) NP-40, 0.05% (w/v) sodium deoxycholate). The washed cell pellet was spun down again and resuspended in fresh Buffer S (50 mM HEPES pH 7.4, 7 M urea, 5% (v/v) glycerol). The cell pellet was solubilized by shaking at room temperature for 60 min. Insoluble debris was spun down at 38,000 g at 4 °C for 30 min. The supernatant was transferred into a new Falcon tube, mixed with pre-equilibrated TALON resin, and incubated with rotation at 4 °C for 2 h. The slurry was transferred to a gravity flow column, and the resin was washed with 2 column volumes of ice-cold Buffer PC500 (50 mM sodium phosphate pH 7.4, 5% glycerol, 500 mM KCl, 0.05% NP-40, 0.2 mM PMSF, freshly added 7 M urea), 3 column volumes of ice-cold Buffer PC100 (50 mM sodium phosphate pH 7.4, 5% glycerol, 100 mM KCl, 0.05% NP-40, 0.2 mM PMSF, 7 M urea), 3 column volumes of PC100 containing 5 mM imidazole and 2 column volumes of PC100 with 15 mM imidazole. Proteins were eluted with 5 column volumes of elution buffer (PC100 with 300 mM imidazole). Successive buffer exchange was performed with buffer PC100 with 0.1% (v/v) NP-40 that contains 4 M, 2 M, 1 M, 0.5 M, and no urea.

To get rid of MYC homodimers, the concentrated and buffer exchanged sample was further purified by gel filtration on a Superose 6 Increase 10/300 column in NMR buffer using the AKTA Purifier system (Cytiva). The protein was aliquoted and stored at -80 °C.

To produce ^15^N-labeled 6×His-tagged MYC IDR, bacteria were cultured in M9 media (42 mM Na_2_HPO_4_, 22 mM KH_2_PO_4_, 8.6 mM NaCl, 18.6 mM ^15^NH_4_Cl, 0.4% (w/v) glucose, 2 mM MgSO_4_, 0.1 mM CaCl_2_), induced and purified using the same protocol as described above.

### Microscale thermophoresis (MST)

The affinity between RNA and protein and between proteins was measured by MST on a Monolith NT.115 (NanoTemper, Germany). Replicates represent individual measurements performed on different days using proteins from the same prep, except for measurements involving 6×His-MBP-GFP-SNIP1-Step-tag-II, 6×His-SMAD4 and 6×His-MYC-Step-tag-II which include two different protein preps.

#### RNA-protein interactions

To measure RNA-protein interactions, we used 50 nM of 6×His-MBP-GFP-30-mers and titrated increasing concentrations of the indicated RNAs. The following assay buffer was used: 25 mM Tris (pH 7.4), 75 mM NaCl, 125 mM KCl, 0.1 mg/ml BSA, 2 mM DTT, 2.5 mM MgCl_2_, 1.25 mM EDTA, 0.01% Tween-20, supplemented with 0.2 U SUPERase•In (Invitrogen).

#### Protein-protein interactions

To measure protein-protein interactions, 50 nM of 6×His-MBP-GFP-MAX-Step-tag-II or 6×His-MBP-GFP-SNIP1-Step-tag-II was used and increasing concentrations of 6×His-tagged proteins (MYC or SMAD4) were added. The experiment was performed in the absence or presence of a constant concentration (40 or 200 nM) of the indicated RNAs in all tubes. The following assay buffer was used: 25 mM Tris (pH 7.4), 150 mM NaCl, 0.1 mg/ml BSA, 2 mM DTT, 5 mM MgCl_2_, 2.5 mM EDTA, 0.01% Tween-20, and 0.2 U SUPERase•In.

PCR tubes were filled with a 1:2 dilution of the titrant, and an equal volume of 100 nM fluorescent molecules was added. The mixtures were incubated at room temperature for 10 min and transferred to 16 standard capillaries (Nanotemper, Germany). Data were acquired using medium MST power, and the blue filter was applied for experiments with GFP-fusion proteins.

#### Calculation of dissociation constant (Kd) from MST measurement

Normalized fluorescence (ΔFnorm) was calculated using NanoTemper Analysis 3. Each individual value was divided by the maximum value of each experiment to obtain “Δ normalized fluorescence”, exported to GraphPad Prism 9 software, and plotted as a function of the concentration of unlabeled titrants where the X axis uses a logarithmic scale. The Kd values were determined by fitting the data from at least two replicates with a one-site specific binding equation in Prism. The binding curve was fitted using the least-squares regression method, and each replicate value is considered as an individual point. The best-fit values for Kd are reported, together with the standard error of fitting. N.A. (not applicable) was reported when the data did not converge when fitted to the one-site specific binding model.

### Knockdown and qPCR experiments

Stable cell lines were generated for shRNA-mediated knockdown experiments. Specifically, pLKO.1/pdR8.2/VSV-G plasmids were co-transfected into HEK293T cells for lentiviral packaging. 48 hours post-transfection, the virus was harvested, aliquoted, and stored at -80 °C or at 4 °C for less than a week. 100 μl virus was used per 12 well to infect U2OS cells. 24 hours after transfection, the virus-containing media was removed, and fresh media was added. Puromycin was added to the medium the next day with a final concentration of 2 μg/ml. cDNA constructs containing GFP-MYC-U were transfected at day 3 after viral transduction of shRNAs.

RNA was isolated from cells by using TRI Reagent (Invitrogen) following the manufacturer’s protocol. RNA was reverse transcribed using qScript(tm) cDNA SuperMix (Quanta Bioscience). cDNA was diluted 2× and used for real-time PCR with gene-specific primers in the presence of PowerUp SYBR Green Master Mix (Life Technologies) by QuantStudio 6 Real-Time PCR system (Applied Biosystems). *ACTB* expression was used as normalization control. Primer sequences are listed in Supplementary Table 3.

### Cotranslational RNA immunoprecipitation (RIP)

RIP experiments were performed using a modified protocol from prior reports^28,29^. HeLa cells were transfected with the indicated constructs and 24 h post-transfection, cells were treated with cycloheximide (100 μg/ml; Sigma Aldrich) or puromycin (50 μg/ml; MP biomedicals) for 15 or 30 min, respectively. The cells were washed twice with PBS and scraped in 500 μl RIP lysis buffer (20 mM HEPES pH 7.5, 150 mM KCl, 10 mM MgCl_2_, 0.5% (v/v) NP-40) supplemented with 40 U/ml SUPERase•In RNase Inhibitor (Invitrogen), EDTA-free protease inhibitor (Roche) and cycloheximide or puromycin.

The crude extracts were incubated on ice for 40 min and sonicated on ice for 1 min (1s on, 2s off). The extracts were cleared by centrifugation at max speed for 15 min, and 10% of the input was saved for RNA extraction. 10 μl of equilibrated GFP-Trap beads was added to the extract and incubated at 4 °C for 2 h. The beads were washed 4 times for 10 min total with high salt wash buffer (20 mM HEPES pH 7.5, 350 mM KCl, 10 mM MgCl_2_, 0.1% (v/v) NP-40) supplemented with SUPERase•In RNase Inhibitor and cycloheximide or puromycin. RNA was extracted from each input and bead sample using TRI Reagent. 1.5 μl of purified RIP-RNA and input RNA samples were used for cDNA synthesis. Enrichment relative to input RNA was calculated using the formula 100 × 2^[(Cp (Input) – 4.907) – Cp (IP)]^ and expressed as “% input RNA”.

### NMR measurement of MYC in the absence and presence of RNAs

All NMR experiments were performed at 298 K using a Bruker Avance-III spectrometer operating at a Larmor Frequency of 700 (16.4 T) and 900 (21 T) MHz equipped with a 5 mm triple resonance (H-C/N-D) cryoprobe. MYC IDR concentration was maintained between 50-75 μM to limit homodimerization. Both free and bound MYC IDR proteins were analyzed using identical experimental conditions and the same protein concentration. The sample for ^1^H-^15^N HSQC spectra of the MYC IDR containing the RNA (*HSPA1B* 3′UTR) was prepared by first diluting stock protein and RNA solutions by 10-20-fold, mixed in of 1:1 molar ratio and concentrated using a centrifugal device with 3 kDa cutoff (Pall) at 4 °C. All experiments were performed in 50 mM sodium phosphate pH 7.4, 100 mM NaCl, 1 mM EDTA, 2.5 mM MgCl_2_, 5 mM DTT, and 5% glycerol. The time-domain data was processed with NMRpipe^54^ and analyzed with Computer-Aided Resonance Assignment (CARA) software.

## Data analysis

### Reanalysis of previously published datasets

#### Compartment bias of mRNAs encoding MYC interactors

MYC interactors were obtained from Tu et al., (2015)^8^ and intersected with information on subcytoplasmic mRNA localization in TGs (TG+) or the cytosol (CY+)^6^.

#### Normalized ensemble diversity (NED) values

NED values were obtained from Ma et al., (2021)^11^ and intersected with information on mRNAs enriched in TGs (TG+) or the cytosol (CY+)^6^.

#### Length and number of IDRs

IUPRED2A (https://iupred2a.elte.hu/) was used to determine IDRs. An IDR was counted if at least 30 uninterrupted amino acids had a score greater than 0.5. The number of all IDRs in a protein was calculated. In addition, the total number of amino acids in these IDRs was determined and intersected with information on subcytoplasmic mRNA localization in TGs (TG+) or the cytosol (CY+)^6^.

### Identification of putative RBH domains and IDR α-helices

UniProt IDs were selected using reviewed Swiss-Prot, Popular Organism Human and protein existence at the protein level, resulting in 16236 proteins. UniProt IDs were downloaded in JSON format for postprocessing. All corresponding AlphaFold2 (AF2) structures were also downloaded. 86 of these structures were excluded as the sequence length in the AF2 structure did not match the sequence length in the UniProt database. The final analysis was carried out on 16150 protein structures.

We used a Python interface for DSSP-based structure prediction^55^ and an adaptation of the DSSP parser in Biobox^56^ to assign all secondary structure elements to each AF2 structure. We retain all α-helices of length 7 amino acids or larger from the secondary structure assignment. Any helix that has an overlap of four amino acids or longer with any known domain as listed under features and domains in the UniProt file is excluded (February 2023). Additionally, regions that fall under Leucine-Zipper, bHLH, DNA-binding region, and PUM-HD, HEAT, or ARM repeats are excluded. 13277 proteins contain at least one α-helix that does not overlap a known domain. However, this number overestimates the true number of α-helices outside of folded domains as many domains are not yet annotated.

#### Serine-rich RBH

Among all α-helices outside of known folded domains, a serine-rich RBH is defined as an α-helix with at least two serines in the five adjacent amino acids, which can be located up- or downstream of the helix. With this definition, we identified 16319 serine-rich RBH domains present in 7921 proteins in the UniProt proteome (Supplementary Table 2). Data and scripts required to rerun the analysis can be found at: https://github.com/meyresearch/alpha_fold_secondary_structure.

#### Amino acid enrichment in the helix-flanking regions

The two groups of α-helices were intersected with information on subcytoplasmic mRNA localization in TGs or the cytosol^6^. All non-membrane proteins expressed in HEK293T cells (*N* = 7015) were analyzed with respect to amino acid enrichment in the helix-flanking regions. We compared 6345 serine-rich RBH domains with 42039 non-serine-rich helices.

### Quantification and statistical analysis

Statistical parameters are reported in the figures and figure legends. Statistical significance is indicated by asterisks: *, *P* < 0.05; **, *P* < 0.01; ***, *P* < 0.001; ****, *P* < 0.0001. Unless otherwise mentioned, a two-sided t-test or two-sided Mann-Whitney test was performed to analyze statistical significance. Analysis was performed in Excel or by GraphPad Prism 9. The amino acid composition of 5-mer sequences that are adjacent to the α-helix within IDRs was obtained by the Biostrings package in R^57^. The chi-square test was performed to identify significantly enriched amino acids adjacent to serine-rich versus non serine-rich helices, for which serine residues are excluded from the analysis.

## Data and code availability

The code to identify α-helices from AlphaFold is available on github (https://github.com/meyresearch/alpha_fold_secondary_structure). All predicted serine-rich RBH domains that are located outside of known folded domains (February 2023) are listed in Supplementary Table 2.

## Figure Legends

**ED Figure 1. 3′UTRs determine biased mRNA localization to TGs or to the cytosol**.

**a**, cDNA constructs for RNA-FISH experiment shown in **b-f**. Yellow marks indicate AU-rich elements (AUUUA) in the 3′UTRs.

**b**, RNA-FISH (cyan) against GFP after transfection of GFP-MYC-U into HeLa cells. BFP-TIS11B (magenta) was co-transfected to visualize TGs. The white dotted lines demarcate the nucleus and the cell boundaries, respectively. Representative images are shown. Right: line profiles of fluorescence intensities. Pearson’s correlation coefficients (R) of the fluorescence intensities were calculated. Quantification of additional cells is shown in **d**.

**c**, As in **b**, but after transfection of GFP-MYC-NU.

**d**, Pearson’s correlation coefficients of fluorescence intensities of BFP-TIS11B and the indicated mRNAs. *N* = 35-43 individual cells were analyzed for each construct. Mann-Whitney test, MYC-U vs. MYC-NU, ****, *P* = E-11; SNIP1-U vs. SNIP-NU, ****, *P* = 7E-17.

**e**, As in **b**, but after transfection of GFP-SNIP1-U.

**f**, As in **b**, but after transfection of GFP-SNIP1-NU.

**g**, The fraction of mRNA transcripts shown as mean ± std encoding MYC and its interaction partners^8^ that localize to TIS granules (yellow) or the cytosol (grey) was obtained from Horste et al. (2022)^6^. Among the MYC interactors, 13 are enriched in TGs (TG+), 16 are enriched in the cytosol (CY+), and 39 have an unbiased (UB) localization pattern. See Supplementary Table 1 for values. The fraction of TG-enriched mRNA transcripts differs significantly among the three groups. Kruskal-Wallis test, ****, *P* = 4.0E-11.

**h**, Western blot showing TIS11B protein expression in a doxycycline inducible CRISPR/Cas9 HeLa cell line expressing either gRNAs targeting *TIS11B* (iKO) or non-targeting controls (Ctrl). Shown is a time course of doxycycline treatment. d, day. GAPDH was used as loading control.

**ED Figure 2. *In vitro* reconstitution of TG-dependent MYC protein complexes**.

**a**, SDS-PAGE for recombinant proteins used for MST and NMR. The MBP-tag is included at the N-terminus to enhance solubility. 6xHis-tag and Strep-Tag II are fused to the N- and C-terminus, respectively, to facilitate purification. CDS, coding sequence.

**b**, Denaturing agarose gel showing the *in vitro*-transcribed RNAs used in MST experiments. Kb, kilobases.

**c**, Mean ± std of MST measurement of the SNIP1-MYC interaction *in vitro* in the absence or presence of 200 nM *DNAJB1* 3′UTR.

**d**, Mean ± std of MST measurement of the MAX-MYC interaction *in vitro* in the presence of 200 nM *HSPA1B* 3′UTR.

**e**, As in **c**, but 200 nM of *MYC* 3′ UTR was used.

**f**, The normalized ensemble diversity (NED) values are shown for 3′UTRs of mRNAs enriched in TGs (TG+, *N =* 1246) or enriched in the cytosol (CY+, *N =* 1481)^6^. NED is a measure that predicts the structural plasticity of an RNA^11^. Mann-Whitney test, *P* = 5.3E-15.

**g**, Table showing the 3′UTR NED values, the percentile of the NED values among mRNAs expressed in HEK293T cells, and the binding affinity of MYC to SNIP1 in the presence of the indicated RNAs. NED, normalized ensemble diversity, which is a measure of RNA structural plasticity.

**ED Figure 3. An α-helix in a serine-rich sequence context is a new RBD within IDRs of transcription factors**.

**a**, Multiple sequence alignment of the MYC IDR from various species. α-helices predicted by AlphaFold are indicated by black boxes. MYC boxes are colored in blue, and yellow squares indicate conserved serine residues adjacent to α-helices. Newly identified RBH are shown.

**b**, Positive and negative charges per amino acid residue (F+ and F-) for the MYC IDR are shown. According to the Das-Pappu diagram^19^, the MYC IDR is considered collapsed. The MYC IDR contains 26 negatively charged and 12 positively charged amino acids.

**c**, AlphaFold prediction of MYC protein structure. Color code as in Fig. 2a. Previously annotated functional domains are colored in dark blue, while serine-rich RBH domains identified in this study are highlighted in yellow.

**d**, SDS-PAGE for recombinant proteins used for RNA affinity measurement by MST.

**e-i**, Mean ± std of MST measurement of the affinity between the indicated GFP-tagged 30-mer peptide and the *HSPA1B* 3′UTR RNA is shown.

**j**, AlphaFold prediction of SNIP1 protein structure. The serine-rich RBH domain is highlighted in yellow.

**k-m**, Mean ± std of MST measurement of the affinity between the indicated GFP-tagged 30-mer peptide and the *HSPA1B* 3′UTR RNA is shown.

**n**, Mean ± std MST measurement of the affinity between GFP-tagged HuR RRM1/2 and a known HuR target RNA (*MYC* 3′UTR) is shown.

**ED Figure 4. *SNIP1* knockdown efficiency**.

*SNIP1* mRNA expression was measured by RT-qPCR in the samples shown in Fig. 3d and was normalized to *ACTB*. Average ± SEM of *N* = 6 biological replicates is shown. Mann-Whitney test, **, *P* = 0.003.

**ED Figure 5. Serine-rich RBH domains in IDRs are widespread**.

**a**, The number of IDRs is shown for proteins encoded by mRNAs enriched in TGs (TG+, *N =* 1246) or enriched in the cytosol (CY+, *N =* 1481)^6^. Only IDRs with ≥ 30 amino acids were included. Mann-Whitney test, *P* = 1.4E-16.

**b**, Schematic of protein domains of CycT1, PIM1, SMAD4, and RBL1.

**c**, Examples of serine-rich RBH domains that bind to the *HSPA1B* 3′UTR. MST data are shown in ED Fig. 3h-i and 5f-h. The sequence of the helix is underlined and the serines in the helix flanking regions are highlighted.

**d**, Examples of IDR α-helices that lack serine-rich flanking regions and do not bind to the *HSPA1B* 3′UTR. The dots indicate additional amino acids in the helices that are not shown. MST data are shown in ED Fig. 5i-m.

**e**, SDS-PAGE for recombinant proteins used for RNA affinity measurement by MST.

**f-m**, Mean ± std of MST measurement of the affinity between the indicated GFP-tagged 30-mer peptide and the *HSPA1B* 3′UTR RNA is shown.

**n**, SDS-PAGE for recombinant SMAD4 protein used for MST.

## Supplementary Tables

**Supplementary Table 1. Fraction of transcripts that localize to TIS granules or the cytosol is shown for mRNAs that encode MYC protein interactors**.

TG, TIS granule; CY, cytosol; UB, unbiased cytoplasmic mRNA localization. Shown is the fraction of mRNA transcripts that localize to the indicated compartments.

**Supplementary Table 2. Serine-rich RBH predicted from AlphaFold**.

Shown are all predicted serine-rich RBH domains from the human UniProt proteome that are located outside of known folded domains. Shown are UniProt IDs, gene names, nucleotide sequence IDs, the position of the predicted α-helices, their sequences as well as their up- and downstream sequences.

**Supplementary Table 3. Primer sequences and other sequences used**.

Shown are sequences of all synthetic oligos used in this study, including primers used for cloning and for generating DNA templates for *in vitro* transcription, qPCR primers, synthetic DNA used to produce the gRNA constructs for generating iKO cell lines, and 30-mer peptides used as GFP-fusions in MST.

