## Supplemental Figures for "mRNA interactions with disordered regions control protein activity"

Luo et al., ED Figure 1

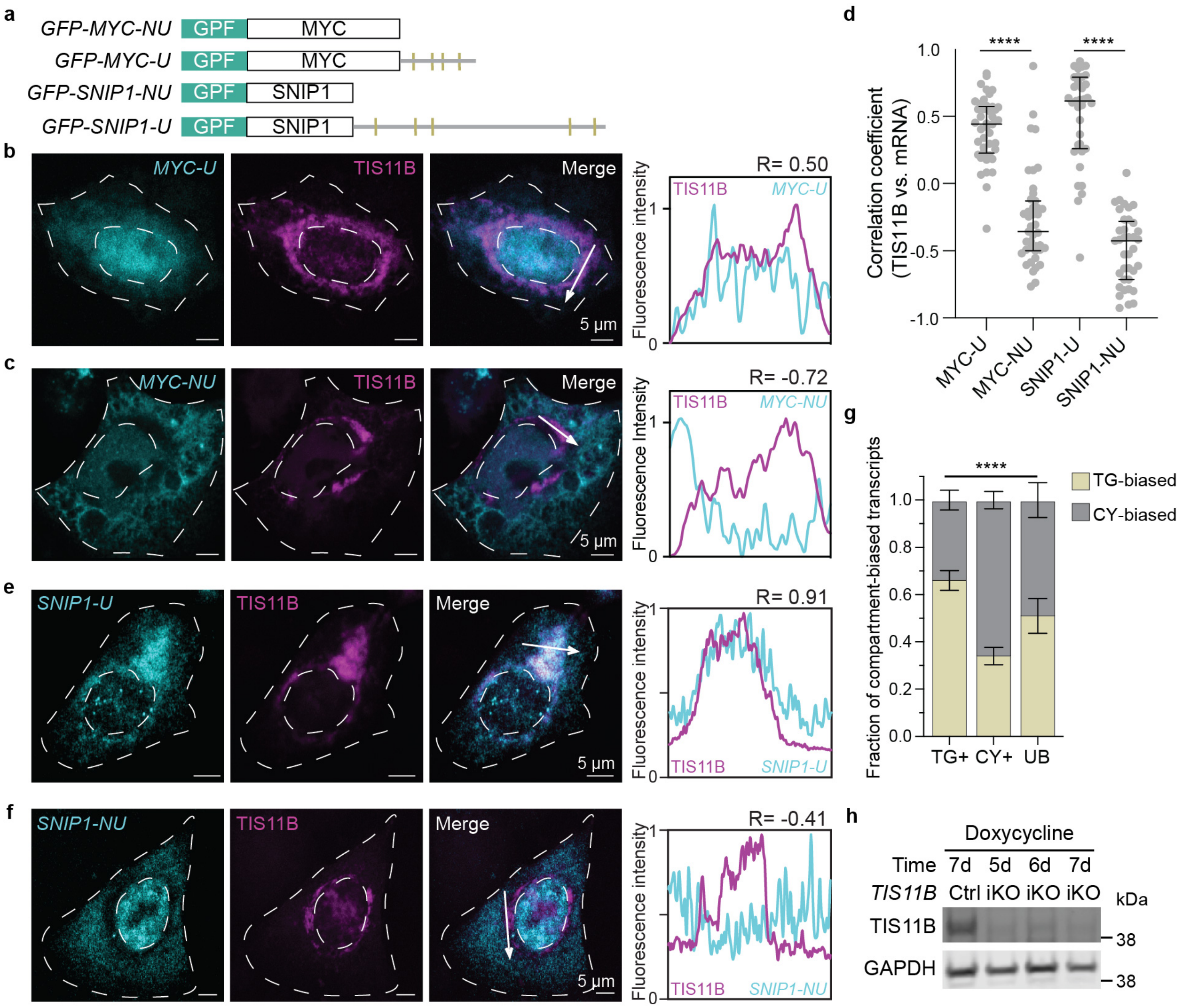

**ED Figure 1. 3′UTRs determine biased mRNA localization to TGs or to the cytosol**.

**a,** cDNA constructs for RNA-FISH experiment shown in **b-f**. Yellow marks indicate AU-rich elements (AUUUA) in the 3′UTRs.

**b,** RNA-FISH (cyan) against GFP after transfection of GFP-MYC-U into HeLa cells. BFP-TIS11B (magenta) was co-transfected to visualize TGs. The white dotted lines demarcate the nucleus and the cell boundaries, respectively. Representative images are shown. Right: line profiles of fluorescence intensities. Pearson's correlation coefficients (R) of the fluorescence intensities were calculated. Quantification of additional cells is shown in **d**.

**c,** As in **b**, but after transfection of GFP-MYC-NU.

**d,** Pearson's correlation coefficients of fluorescence intensities of BFP-TIS11B and the indicated mRNAs. *N* = 35-43 individual cells were analyzed for each construct. Mann-Whitney test, MYC-U vs. MYC-NU, ****, *P* = E-11; SNIP1-U vs. SNIP-NU, ****, *P* = 7E-17.

**e,** As in **b**, but after transfection of GFP-SNIP1-U.

**f,** As in **b**, but after transfection of GFP-SNIP1-NU.

**g,** The fraction of mRNA transcripts shown as mean ± std encoding MYC and its interaction partners[8] that localize to TIS granules (yellow) or the cytosol (grey) was obtained from Horste et al. (2022)[6]. Among the MYC interactors, 13 are enriched in TGs (TG+), 16 are enriched in the cytosol (CY+), and 39 have an unbiased (UB) localization pattern. See Supplementary Table 1 for values. The fraction of TG-enriched mRNA transcripts differs significantly among the three groups. Kruskal-Wallis test, ****, *P* = 4.0E-11.

**h,** Western blot showing TIS11B protein expression in a doxycycline inducible CRISPR/Cas9 HeLa cell line expressing either gRNAs targeting *TIS11B* (iKO) or non-targeting controls (Ctrl). Shown is a time course of doxycycline treatment. d, day. GAPDH was used as loading control.

Luo et al., ED Figure 2

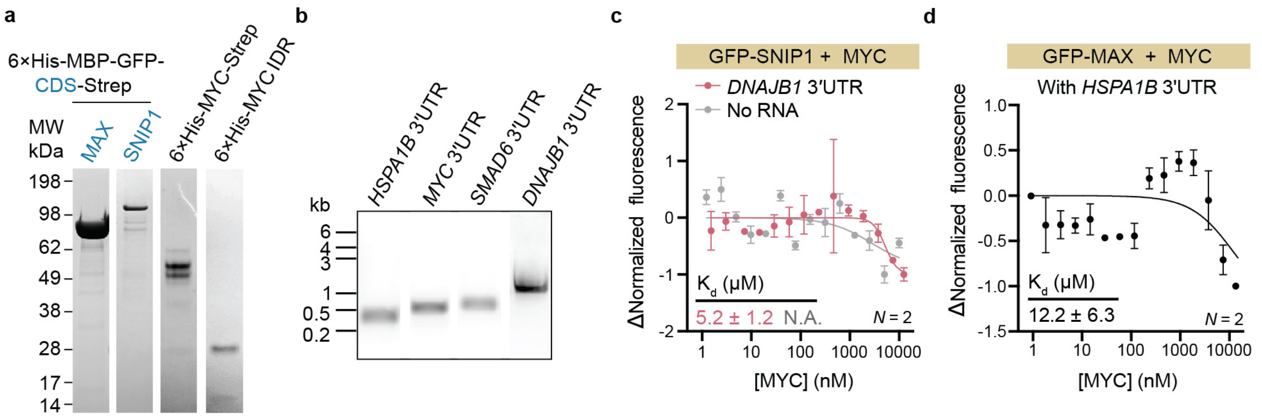


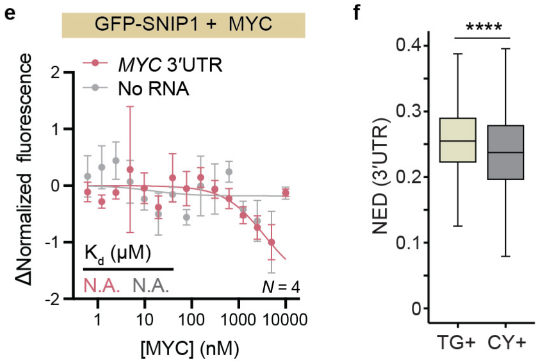


g

| 3′UTR | NED | Percentile | Kd (SNIP1-MYC) (μM) |
|---|---|---|---|
| *HSPA1B* | 0.36 | 2 | 1.8 ± 0.4 |
| *DNAJB1* | 0.28 | 28 | 5.2 ± 1.2 |
| *MYC* | 0.22 | 67 | N.A. |
| *SMAD6* | 0.21 | 72 | N.A. |

**ED Figure 2. *In vitro* reconstitution of TG-dependent MYC protein complexes.**

**a,** SDS-PAGE for recombinant proteins used for MST and NMR. The MBP-tag is included at the N-terminus to enhance solubility. 6xHis-tag and Strep-Tag II are fused to the N- and C-terminus, respectively, to facilitate purification. CDS, coding sequence.

**b,** Denaturing agarose gel showing the *in vitro*-transcribed RNAs used in MST experiments. Kb, kilobases.

**c**, Mean ± std of MST measurement of the SNIP1-MYC interaction *in vitro* in the absence or presence of 200 nM *DNAJB1* 3′UTR.

**d**, Mean ± std of MST measurement of the MAX-MYC interaction *in vitro* in the presence of 200 nM *HSPA1B* 3′UTR.

**e,** As in **c**, but 200 nM of *MYC* 3′ UTR was used.

**f,** The normalized ensemble diversity (NED) values are shown for 3′UTRs of mRNAs enriched in TGs (TG+, *N* = 1246) or enriched in the cytosol (CY+, *N* = 1481)[6]. NED is a measure that predicts the structural plasticity of an RNA[11]. Mann-Whitney test, *P* = 5.3E-15.

**g,** Table showing the 3′UTR NED values, the percentile of the NED values among mRNAs expressed in HEK293T cells, and the binding affinity of MYC to SNIP1 in the presence of the indicated RNAs. NED, normalized ensemble diversity, which is a measure of RNA structural plasticity.

Luo et al., ED Figure 3

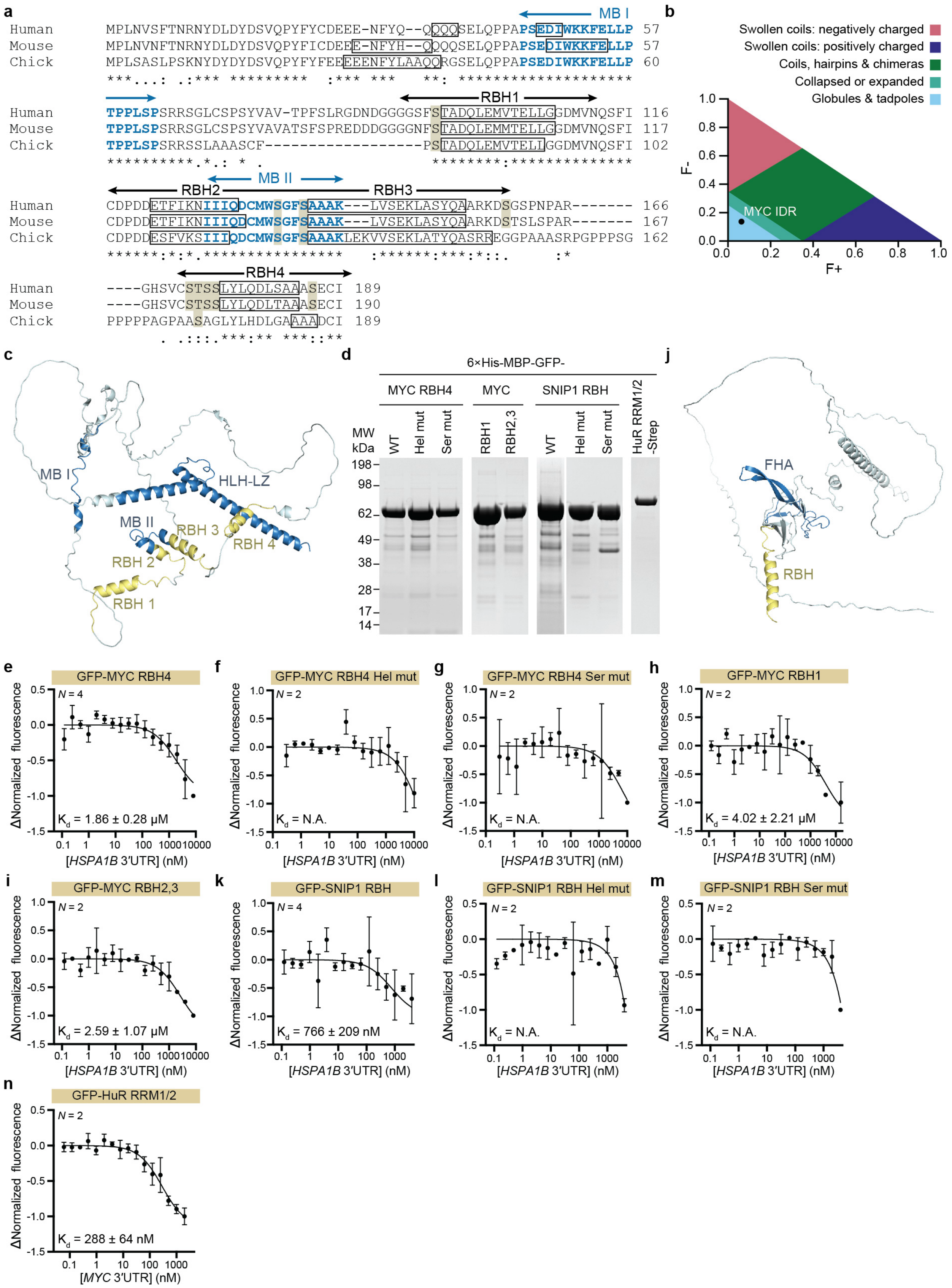

**ED Figure 3. An α-helix in a serine-rich sequence context is a new RBD within IDRs of transcription factors.**

**a,** Multiple sequence alignment of the MYC IDR from various species. α-helices predicted by AlphaFold are indicated by black boxes. MYC boxes are colored in blue, and yellow squares indicate conserved serine residues adjacent to α-helices. Newly identified RBH are shown.

**b,** Positive and negative charges per amino acid residue (F+ and F-) for the MYC IDR are shown. According to the Das-Pappu diagram[19], the MYC IDR is considered collapsed. The MYC IDR contains 26 negatively charged and 12 positively charged amino acids.

**c**, AlphaFold prediction of MYC protein structure. Color code as in Fig. 2a. Previously annotated functional domains are colored in dark blue, while serine-rich RBH domains identified in this study are highlighted in yellow.

**d,** SDS-PAGE for recombinant proteins used for RNA affinity measurement by MST.

**e-i,** Mean ± std of MST measurement of the affinity between the indicated GFP-tagged 30-mer peptide and the *HSPA1B* 3′UTR RNA is shown.

**j**, AlphaFold prediction of SNIP1 protein structure. The serine-rich RBH domain is highlighted in yellow.

**k-m,** Mean ± std of MST measurement of the affinity between the indicated GFP-tagged 30-mer peptide and the *HSPA1B* 3′UTR RNA is shown.

**n**, Mean ± std MST measurement of the affinity between GFP-tagged HuR RRM1/2 and a known HuR target RNA (*MYC* 3′UTR) is shown.

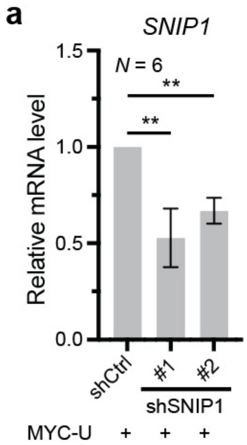


**ED Figure 4. *SNIP1* knockdown efficiency.**

*SNIP1* mRNA expression was measured by RT-qPCR in the samples shown in Fig. 3d and was normalized to *ACTB*. Average ± SEM of $N$ = 6 biological replicates is shown. Mann-Whitney test, **, $P$ = 0.003.

Luo et al., ED Figure 5

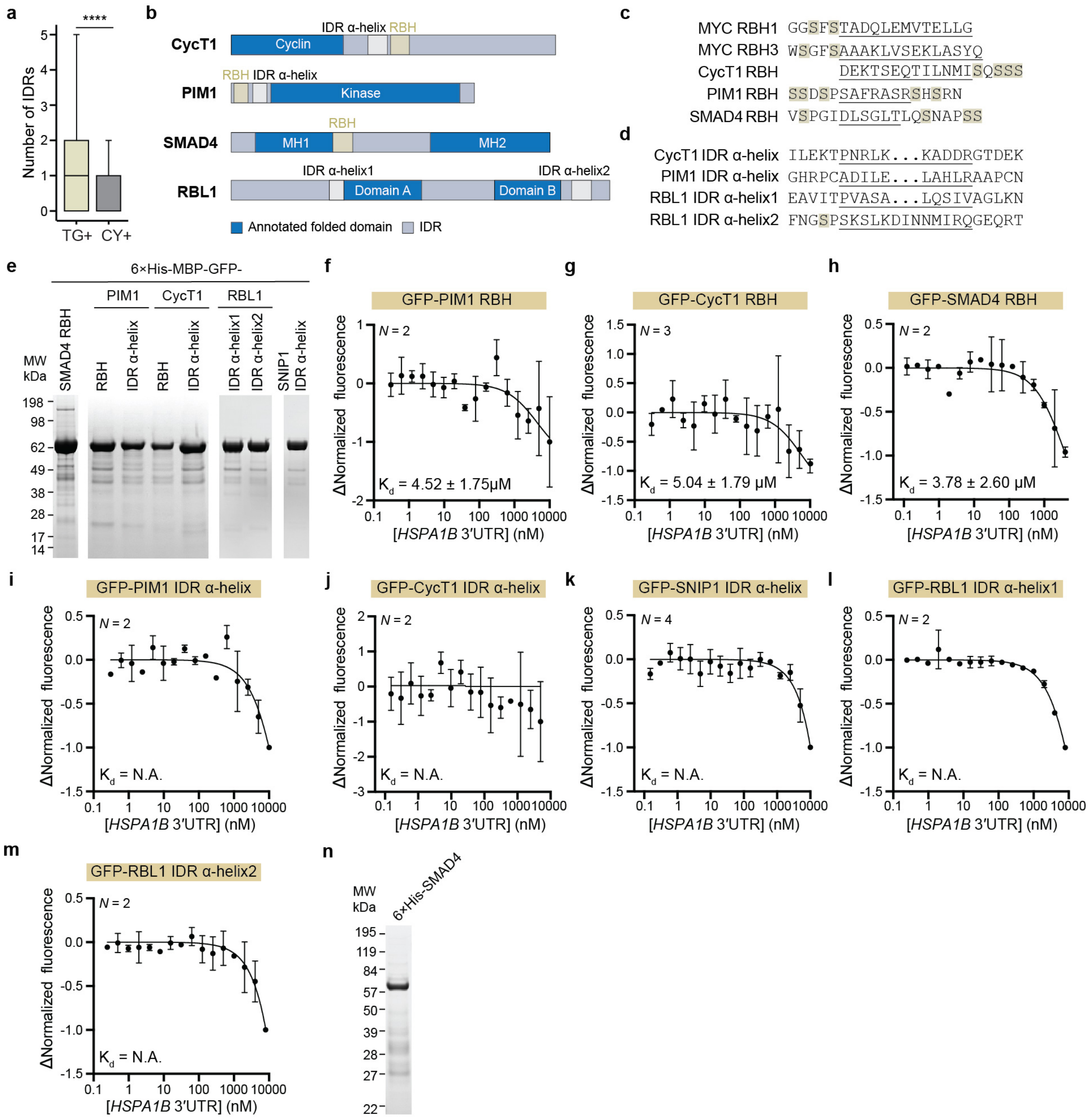

**ED Figure 5. Serine-rich RBH domains in IDRs are widespread**.

**a,** The number of IDRs is shown for proteins encoded by mRNAs enriched in TGs (TG+, *N* = 1246) or enriched in the cytosol (CY+, *N* = 1481)[6].Only IDRs with ≥ 30 amino acids were included. Mann-Whitney test, *P* = 1.4E-16.

**b,** Schematic of protein domains of CycT1, PIM1, SMAD4, and RBL1.

**c,** Examples of serine-rich RBH domains that bind to the *HSPA1B* 3′UTR. MST data are shown in ED Fig. 3h-i and 5f-h. The sequence of the helix is underlined and the serines in the helix flanking regions are highlighted.

**d,** Examples of IDR α-helices that lack serine-rich flanking regions and do not bind to the *HSPA1B* 3′UTR. The dots indicate additional amino acids in the helices that are not shown. MST data are shown in ED Fig. 5i-m.

**e,** SDS-PAGE for recombinant proteins used for RNA affinity measurement by MST.

**f-m**, Mean ± std of MST measurement of the affinity between the indicated GFP-tagged 30-mer peptide and the *HSPA1B* 3′UTR RNA is shown.

**n,** SDS-PAGE for recombinant SMAD4 protein used for MST.
